# Revealing nonlinear neural decoding by analyzing choices

**DOI:** 10.1101/332353

**Authors:** Qianli Yang, Edgar Walker, R. James Cotton, Andreas S. Tolias, Xaq Pitkow

## Abstract

Sensory data about most natural task-relevant variables are entangled with task-irrelevant nuisance variables. The neurons that encode these relevant signals typically constitute a nonlinear population code. Here we present a theoretical framework for quantifying how the brain uses or decodes its nonlinear information. Our theory obeys fundamental mathematical limitations on information content inherited from the sensory periphery, identifying redundant codes when there are many more cortical neurons than primary sensory neurons. The theory predicts that if the brain uses its nonlinear population codes optimally, then more informative patterns should be more correlated with choices. More specifically, the theory predicts a simple, easily computed quantitative relationship between fluctuating neural activity and behavioral choices that reveals the decoding efficiency. We analyze recordings from primary visual cortex of monkeys discriminating the distribution from which oriented stimuli were drawn, and find these data are consistent with the hypothesis of near-optimal nonlinear decoding.

## 1 Introduction

How does an animal use, or ‘decode’, the information represented in its brain? When the average responses of some neurons are well-tuned to a stimulus of interest, this is straightforward. In binary discrimination tasks, for example, a choice can be reached simply by a linear weighted sum of these tuned neural responses. Yet real neurons are rarely tuned to precisely one variable: variation in multiple stimulus dimensions influence their responses in complex ways. As we show below, when these nuisance variations have nonlinear effects on responses, they can dilute or even abolish the mean tuning to the relevant stimulus. Then the brain cannot simply use linear computation, nor can we understand neural processing using linear models.

A quantitative account of nonlinear neural decoding of sensory stimuli must first express how populations of neurons encode or represent information. Past theories of nonlinear population codes made unsupported assumptions about the variability of these population responses [1, 2], substantially underestimating the redundancy of large cortical populations that are driven by a smaller population of sensory inputs. Here we correct this problem by generalizing information-limiting correlations [3] to nonlinear population codes, providing a more realistic theory of how much sensory information is encoded in the brain.

Just because a neural population encodes information, it does not mean that the brain decodes it all. Here, *encoding* specifies how the neural responses relate to the stimulus input, whereas *decoding* specifies how the neural responses relate to the behavioral output. To understand the brain’s computational strategy we must understand how encoding and decoding are related, *i.e.* how the brain uses the information it has. These are distinct processes, so the brain could encode a stimulus well while decoding it poorly, or vice-versa.

We provides a simple way of testing the hypothesis that the brain’s decoding strategy is efficient, using a simple, novel statistic to assess whether neural response patterns that are informative about the task-relevant sensory input are also informative about the animal’s behavior in the task. This extends past results on linear decoding [4] by relaxing multiple assumptions that are violated when neuronal response statistics beyond the mean are tuned.

These key theoretical advances are developed in general, and applied to simple models to illustrate the approach. Finally, we apply this test to analyze V1 data from macaque monkeys, finding direct experimental evidence for optimal nonlinear decoding.

## 2 Results

### 2.1 Framework for nonlinear computation

#### 2.1.1 A simple example of a nonlinear code

Imagine a simplified model of a visual neuron that includes an oriented edge-detecting linear filter followed by additive noise, with a Gabor receptive field like simple cells in primary visual cortex (Figure 1A). If an edge is presented to this model neuron, different rotation angles will change the overlap, producing a different mean. This neuron is then tuned to orientation.

**Figure 1:**
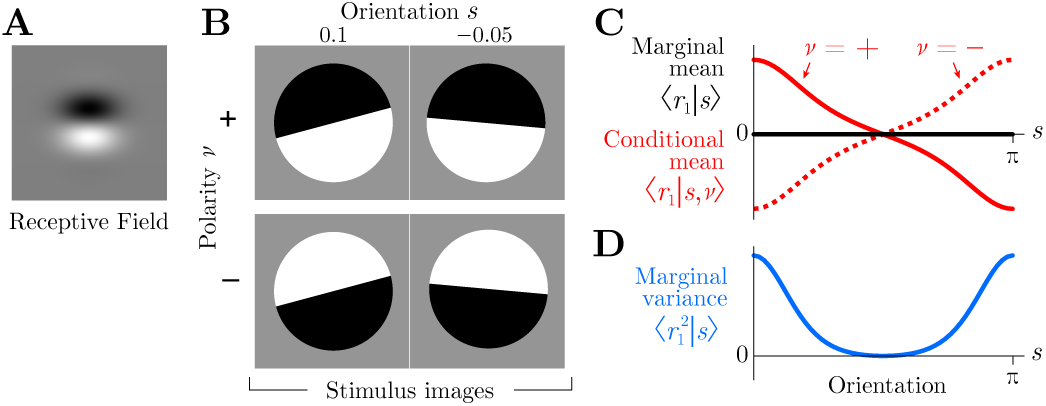
Simple nonlinear code for orientation induced by two polarities. (**A**) Receptive field for a linear neuron. (**B**) Four example images, each with an orientation *s* ∈ [0, *π*) and a polarity *ν* ∈ {−1, +1}. (**C**) The mean response of the linear neuron is tuned to orientation if polarity were specified (conditional mean, red). But when the polarity is unknown and could take either value, the mean response is untuned (marginal mean, black). (**D**) Tuning is recovered by the marginal variance even if the polarity is unknown (blue).

However, when the edge has the opposite polarity, with black and white reversed, then the linear response is reversed also. If the two polarities occur with equal frequency, then the positive and negative responses cancel on average. The mean response of this linear neuron to any given orientation is therefore precisely constant, so the model neuron is untuned.

Notice that stimuli aligned with the neuron’s preferred orientation will generally elicit the highest or lowest response magnitude, depending on polarity. Edges evoking the largest response to one polarity will also evoke the smallest response to its inverse. Thus, even though the mean response of this linear neuron is zero, independent of orientation, the *variance* is tuned.

To estimate the variance, and thereby the orientation itself, the brain can compute the square of the linear responses. This would allow the brain to estimate the orientation independently from polarity. This is consistent with the well-known energy model of complex cells in primary visual cortex, which use squaring nonlinearities to achieve invariance to the polarity of an edge [5].

Generalizing from this example, we identify edge polarity as a ‘nuisance variable’ — a property in the world that alters how task-relevant stimuli appear but is, itself, irrelevant for the current task (here, perceiving orientation). Other examples of nuisance variables include the illuminant for guessing surface color, position for object recognition, expression for face identification, or pitch for speech recognition. Generically, nuisance variables make it hard to extract the task-relevant variables from sense data, which is the central task of perception [6–9]. For example, cells in early visual cortex are not tuned to object identity, since the object could appear at any location and V1 has not yet extracted the complex combinations of features that reveal object type independent of the nuisance variable of position. (Of course, what is a nuisance for one task might be a target variable in another task, and vice versa.)

The prevailing neuroscience view of this disentangling process is deterministic: the output of a complex (often multi-stage) nonlinear function extracts the variables of interest [6, 7, 10]. But can we understand *which* nonlinear function is useful, beyond simply stating what works? A statistical perspective helps here: the brain learns from its history of sensory inputs which statistics of its many sense data are tuned to the task-relevant variable. Good nonlinear computations then compute those statistics. In the orientation estimation task above, the relevant statistic was not the mean, but the variance.

The next sections formalize and develop these ideas to describe redundant nonlinear population codes.

#### 2.1.2 Task, stimuli, neural responses, actions

Our mathematical framework describes a perceptual task, a stimulus with both relevant and irrelevant variables, neural responses, and behavioral choices.

In our task, an agent observes a multidimensional stimulus (*s, ν*) and must act upon one particular relevant aspect of that stimulus, *s*, while ignoring the rest, *ν*. The irrelevant stimulus aspects serve as nuisance variables for the task (*ν* is the Greek letter ‘nu’ and here stands for *nu*isance). Together, these stimulus properties determine a complete sensory input that drives some responses ***r*** in a population of *N* neurons according to the distribution *p*(***r***|*s, ν*).

We consider a feedforward processing chain for the brain, in which the neural responses ***r*** are nonlinearly transformed downstream into other neural responses ***R***(***r***), which in turn are used to create a perceptual estimate of the relevant stimulus *ŝ*:

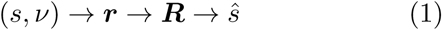

We model the brain’s estimate as a linear function of the downstream responses ***R***. Ultimately these estimates are used to generate an action that the experimenter can observe. We assume that we have recorded activity only from some of the upstream neurons, so we don’t have direct access to ***R***, only a subset of ***r***. Nonetheless we would like to learn something about the downstream computations used in decoding. In this paper we show how to use the statistics of cofluctuations in ***r***, *s*, and *ŝ* to estimate the quality of nonlinear decoding.

We first develop the theory for local or fine-scale estimation tasks: the subject must directly report its estimate *ŝ* for the relevant stimuli near a reference *s*_0_, and we measure performance by the variance of this estimate, 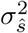. This fine-scale continuous estimation provides the simplest mathematical framing of the problem. In later sections we then generalize the problem to allow for binary discrimination as well as coarse tasks. These binary and coarse discriminations are not conceptually different from fine estimation, but the mathematical details are more complicated.

#### 2.1.3 Signal and noise

The population response, which we take here to be the spike counts of each neuron in a specified time window, reflects both *signal* and *noise*, where signal is the repeatable stimulus-dependent aspects of the response, and noise reflects trial-to-trial variation. Conventionally in neuroscience, the signal is often thought to be the stimulus dependence of the *average* response, *i.e.* the tuning curve ***f*** (*s*) = ∑_***r***_ ***r*** *p*(***r***|*s*) = ⟨***r***|*s*⟩ (angle brackets denote an average over all responses given the condition after the vertical bar). Below we will broaden this conventional definition to allow the signal to include any stimulus-dependent statistical property of the population response.

Noise is the non-repeatable part of the response, characterized by the variation of responses to a fixed stimulus. It is convenient to distinguish *internal* noise from *external* noise. Internal noise is internal to the animal, and is described by response distribution *p*(***r***|*s, ν*) when everything about the stimulus is fixed. This could also include uncontrolled variation in internal states [11–14], like attention, motivation, or wandering thoughts. External noise is variability generated by the external world, or nuisance variables. Whether this should count as ‘noise’ is somewhat contentious. In some instances, most people readily describe external variation as noise, as for a ‘white noise stimulus’ or a random dot kinematogram. In other cases people might be more reticent to label this variability as noise, as for the uncontrolled polarity of an edge (Figure 1) or the lighting of a three-dimensional scene. Regardless of the name, external variability leads to a neural response distribution *p*(***r***|*s*) where only the relevant variables are held fixed. Both types of noise can lead to uncertainty about the true stimulus.

Trial-to-trial variability can of course be correlated across neurons. Neuroscientists often measure two types of second-order correlations: signal correlations and noise correlations [2, 15–22]. Signal correlations measure shared variation in mean responses ***f***(*s*) averaged over the set of stimuli *s*: *ρ*_signal_ = Corr(***f***(*s*)). (Internal) noise correlations measure shared variation that persists even when the stimulus is completely identical, nuisance variables and all: *ρ*_noise_(*s, ν*) = Corr(***r***|*s, ν*).

For multidimensional stimuli, however, these are only two extremes on a spectrum, depending on how many stimulus aspects are fixed across the trials to be averaged. We propose an intermediate type of correlation: *nuisance correlations*. Here we fix the task-relevant stimulus variable(s) *s*, and average over the nuisance variables *ν*: *ρ*_nuisance_(*s*) = Corr(***f*** (*ν*)|*s*). Including both internal and external (nuisance) noise correlations gives Corr (***r***|*s*).

Critically, but confusingly, some so-called ‘noise’ correlations and nuisance correlations actually serve as signals. This happens whenever the statistical pattern of trial-by-trial fluctuations depends on the stimulus, and thus contain information. For example, a stimulus-dependent noise covariance functions as a signal. There would still be true noise, *i.e.* irrelevant trial-to-trial variability that makes the signal uncertain, but it would be relegated to higher-order fluctuations [23] such as the variance of the response covariance (Figure 2D, Table 1). Whether from internal or external noise, stimulus-dependent correlations lead naturally to nonlinear population codes, as we explain below.

**Table 1:**
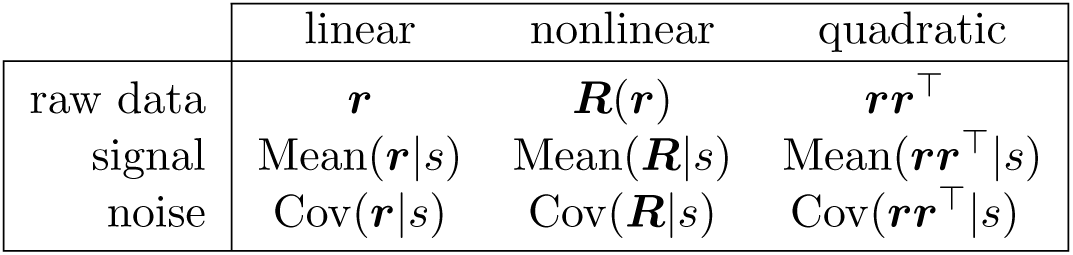
Neural response properties relevant for linear and nonlinear codes. In each case, the brain must estimate the stimulus from a single example of neural data, but the relevant function of that data is linear for linear codes, and nonlinear for nonlinear codes (such as the quadratic example in the last column). The noise and signal can be quantified by the corresponding covariance and stimulus-dependent changes in the corresponding means (*i.e.* the tuning curve slope).

**Figure 2:**
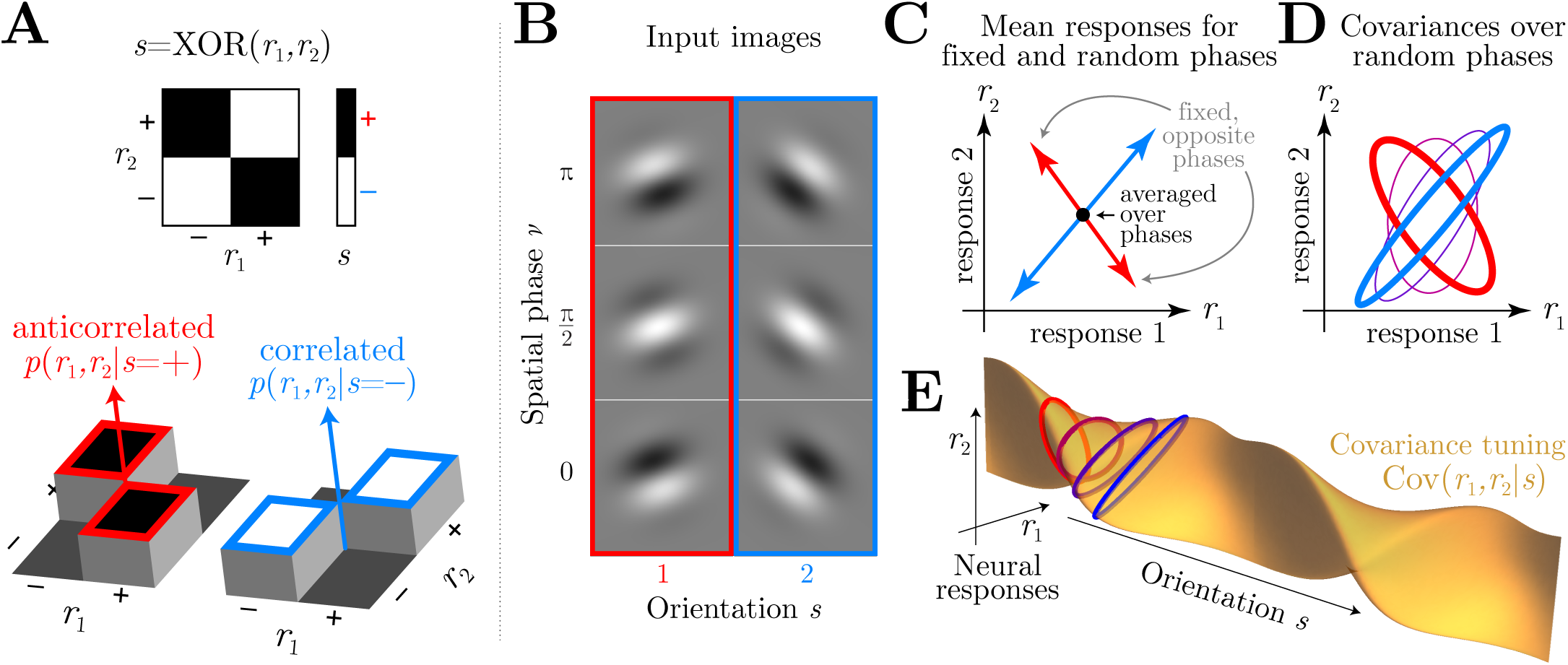
Nonlinear codes. (**A**) Simple example in which a stimulus *s* is the XOR of two neural responses (top). Conditional probabilities *p*(*r*_1_, *r*_2_|*s*) of those responses (bottom) show they are anti-correlated when *s* = +1 (red) and positively correlated when *s* = −1 (blue). This stimulus-dependent correlation between responses creates a nonlinear code. The remaining panels show that a similar stimulus-dependent correlation emerges in orientation discrimination with unknown spatial phase. (**B**) Gabor images with two orientations and three spatial phases. (**C**) Mean responses of linear neurons with Gabor receptive fields are sensitive to orientation when phase is fixed (arrows), but point in different directions for different spatial phases. When phase is an unknown nuisance variable, this mean tuning therefore vanishes (black dot). (**D**) The response covariance Cov(*r*_1_, *r*_2_|*s*) between these linear neurons is tuned to orientation even when averaging over spatial phase. Response covariances for four orientations are depicted by ellipses. (**E**) A continuous view of the covariance tuning to orientation for a pair of neurons.

#### 2.1.4 Nonlinear encoding by neural populations

Most accounts of neural population codes actually address *linear* codes, in which the mean response is tuned to the variable of interest and completely captures all signal about it [3, 24–27]. We call these codes linear because the neural response property needed to best estimate the stimulus near a reference (or even infer the entire likelihood of the stimulus, Supplement S.1.2.2) is a linear function of the response. Linear codes for different variables may arise early in sensory processing, like orientation in V1, or after many stages of computation [6, 9], like for objects in inferotemporal cortex.

If any of the relevant signal can only be extracted using nonlinear functions of the neural responses, then we say that the population code is nonlinear.

It is illuminating to take a statistical view: unlike a linear code, the information is not encoded in mean neural responses but instead by higher-order statistics of responses [1, 2]. These functional and statistical views are naturally linked because estimating higher-order statistics requires nonlinear operations. For instance, information from a stimulus-dependent covariance *Q*(*s*) = ⟨***rr***^T^|*s*⟩ can be decoded by quadratic operations ***R*** = ***rr***^T^ [28–30]. Table 1 compares the relevant neural response properties for linear and nonlinear codes.

A simple example of a nonlinear code is the exclusive-or (XOR) problem. Given the responses of two binary neurons, *r*_1_ and *r*_2_, we would like to decode the value of a task-relevant signal *s* = XOR(*r*_1_, *r*_2_) (Figure 2A). We don’t care about the specific value of *r*_1_ by itself, and in fact *r*_1_ alone tells us nothing about *s*. The same is true for *r*_2_. The usual view on nonlinear computation is that the desired signal can be extracted by applying an XOR or product nonlinearity. However, there is an underlying statistical reason this works: the signal is actually reflected in the trial-by-trial *correlation* between *r*_1_ and *r*_2_: when they are the same then *s* = −1, and when they are opposite then *s* = +1. The correlation, and thus the relevant variable *s*, can be estimated nonlinearly from *r*_1_ and *r*_2_ as *ŝ* = −*r*_1_*r*_2_.

Some experiments have reported stimulus-dependent internal noise correlations that depend on the signal, even for a completely fixed stimulus without any nuisance variation [31–35]. Other experiments have turned up evidence for nonlinear population codes by characterizing the nonlinear selectivity directly [10, 36, 37].

More typically, however, stimulus-dependent correlations arise from external noise, leading to what we call nuisance correlations. In the introduction (Figure 1) we showed a simple orientation estimation example in which fluctuations of an unknown polarity eliminate the orientation tuning of mean responses, relegating the tuning to variances. Figure 2B–E shows a slightly more sophisticated version of this example, where instead of two image polarities, we introduce spatial phase as a continuous nuisance variable. This again eliminates mean tuning, but introduces nuisance covariances that are orientation tuned.

One might object that although the nuisance covariance is tuned to orientation, a subject cannot compute the covariance on a single trial because it does not experience all possible nuisance variables to average over. This objection stems from a conceptual error that conflates the tuning (signal) with the raw sense data (signal+noise). In linear codes, the subject does not have access to the tuned mean response ⟨***r***|*s*⟩ either, just a noisy single-trial version of the mean, namely ***r***. Analogously, the subject does not need access to the tuned covariance, just a noisy single-trial version of the second moments, ***rr***^⊺^ (Table 1). In this simple example, the nuisance variable of spatial phase ensures that quadratic statistics contains relevant information about the orientation, just like complex cells in V1 [5].

### 2.2 Decoding and choice correlations

To study how neural information is used or decoded, past studies have examined whether neurons that are sensitive to sensory inputs also reflect an animal’s behavioral outputs or choices [38–46]. However, this choice-related activity is hard to interpret, because it may reflect decoding of the recorded neurons, or merely correlations between them and other neurons that are decoded instead [47].

In principle, we could discount such indirect relationships with complete recordings of all neural activity. This is currently impractical for most animals, and even if we could record from all neurons simultaneously, we would struggle to acquire enough trials to fully disambiguate how neural activities directly influence behavior.

To understand key principles of neural computation, however, we may not care about all detailed patterns of decoding weights and their underlying synaptic connectivity. Instead we may want to know only certain properties of the brain’s strategies. One important property is the efficiency with which the brain decodes available neural information as it generates an animal’s choices.

Conveniently, testable predictions about choice-related activity can reveal the brain’s decoding efficiency, in the case of linear codes [27]. Next we review these predictions, and then generalize them to nonlinear codes.

#### 2.2.1 Choice correlations predicted for optimal linear decoding

We define ‘choice correlation’ 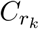 as the correlation coefficient between the response *r*_*k*_ of neuron *k* and the stimulus estimate (which we view as a continuous ‘choice’) *ŝ*, given a fixed stimulus *s*:

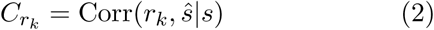

This choice correlation is a conceptually simpler and more convenient measure than the more conventional statistic, ‘choice probability’ [48], but it has almost identical properties (Methods 4.2) [27, 47].

Intuitively, if an animal is decoding its neural information efficiently, then those neurons encoding more information should be more correlated with the choice. Mathematically, one can show that choice correlations indeed have this property when decoding is optimal [27]:

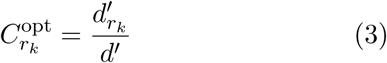

where *d*′ and 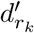 are, respectively, the stimulus discriminability [49] based on the entire population ***r*** or on neuron *k*’s response *r*_*k*_ (Methods 4.2). This relationship holds for a locally optimal linear estimator,

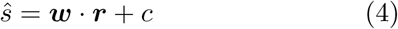

for any stimulus-independent noise correlations, regardless of their structure.

Another way to test for optimal linear decoding would be to measure whether the animal’s behavioral discriminability matches the discriminability for an ideal observer of the neural population response. Yet this approach is not feasible, as it requires one to measure simultaneous responses of many, or even all, relevant neurons. In contrast, the optimality test (Eq. 3) requires measuring only non-simultaneous single neuron responses, which is vastly easier. Neural recordings in the vestibular system are consistent with near-optimal decoding according to this prediction [27].

#### 2.2.2 Nonlinear choice correlations for optimal decoding

However, when nuisance variables wash out the mean tuning of neuronal responses, we may well find that a single neuron has both zero choice correlation and zero information about the stimulus. The optimality test would thus be inconclusive.

This situation is exactly the same one that gives rise to nonlinear codes. A natural generalization of Equation 3 can reveal the quality of neural computation on nonlinear codes. We simply define a ‘*nonlinear* choice correlation’ between the stimulus estimate *ŝ* and nonlinear functions of neural activity ***R***(***r***):

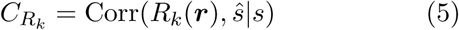

(Methods 4.2), where *R*_*k*_(***r***) is a nonlinear function of the neural responses. If the brain optimally decodes the information encoded in the nonlinear statistics of neural activity, according to the simple nonlinear extension to Eq. 4,

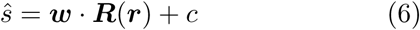

then the nonlinear choice correlation satisfies the equation

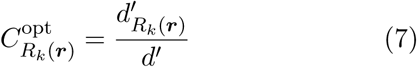

where 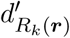 is the stimulus discriminability provided by *R*_*k*_(***r***) (Methods 4.2.1).

As an example of this relationship, we return to the orientation example. Here the response covariance Σ(*s*) = Cov(***r***|*s*) depends on the stimulus, but the mean ***f*** = ⟨***r***|*s*⟩ = ⟨***r***⟩ does not. In this model, optimally decoded neurons would have no linear correlation with behavioral choice. Instead, the choice should be driven by the product of the neural responses, ***R***(***r***) = vec(***rr***^⊺^), where vec(·) is a vectorization that flattens an array into a one-dimensional list of numbers. Such quadratic computation is what the energy model for complex cells is thought to accomplish for phase-invariant orientation coding [5]. Figure 3 shows linear and nonlinear choice correlations for pairs of neurons, defined as 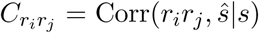. When decoding is linear, linear choice correlations are strong while nonlinear choice correlations are near zero (Figure 3A,B). When the decoding is quadratic, here mediated by an intermediate layer that multiplies pairs of neural activity, the nonlinear choice correlations are strong while the linear ones are insignificant (Figure 3C,D).

**Figure 3:**
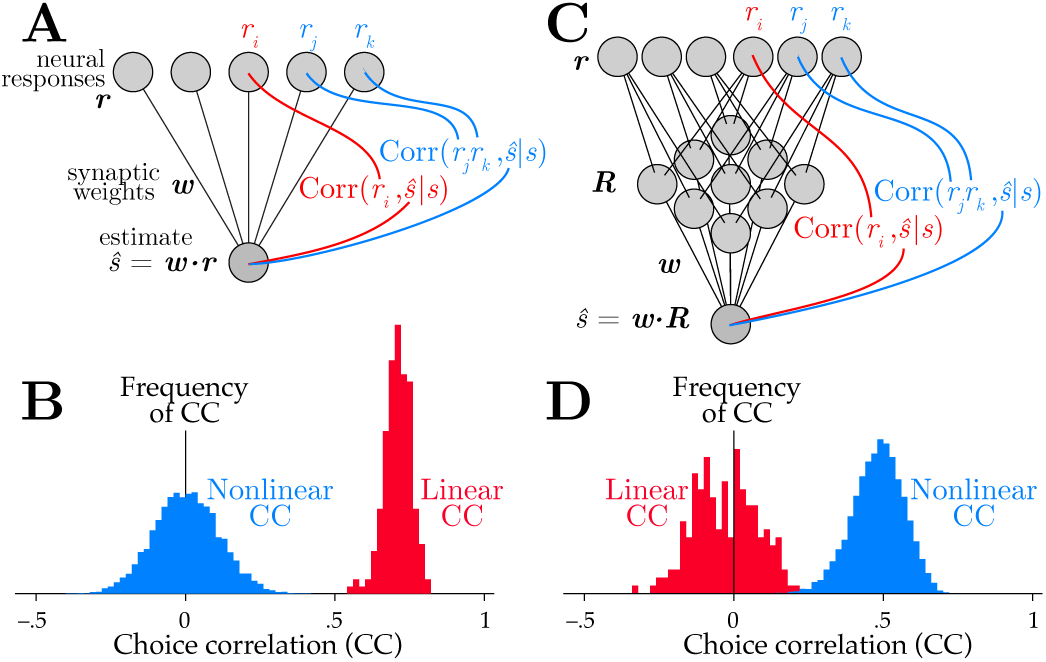
Linear and nonlinear choice correlations successfully distinguish network structure. A linearly decoded population (**A**) produces nonzero linear choice correlations (**B**), while the nonlinear choice correlations are randomly distributed around zero. The situation is reverse for a nonlinear network (**C**), with insignificant linear choice correlations but strong nonlinear ones (**D**). Here the network implements a quadratic nonlinearity, so the relevant choice correlations are quadratic as well, *C*_*jk*_ = Corr(*r*_*j*_*r*_*k*_, *ŝ*|*s*).

#### 2.2.3 Which nonlinearity?

If the brain’s decoder optimally uses all available information, choice correlations will obey the prediction of Eq. 7 even if the specific nonlinearities used by the brain differ from those selected for evaluating choice correlations (Methods 4.2.2). The prediction is valid as long as the brain’s nonlinearity can be expressed as a linear combination of the tested nonlinearities (Methods 4.2.2). Figure 4 shows a situation where information is encoded by linear, quadratic and cubic sufficient statistics of neural responses, while a simulated brain decodes them near-optimally using a generic neural network rather than a set of nonlinearities matched to those sufficient statistics. Despite this mismatch we can successfully identify that the brain is near-optimal by applying Eq. 7, even without knowing details of the simulated brain’s true nonlinear transformations.

**Figure 4:**
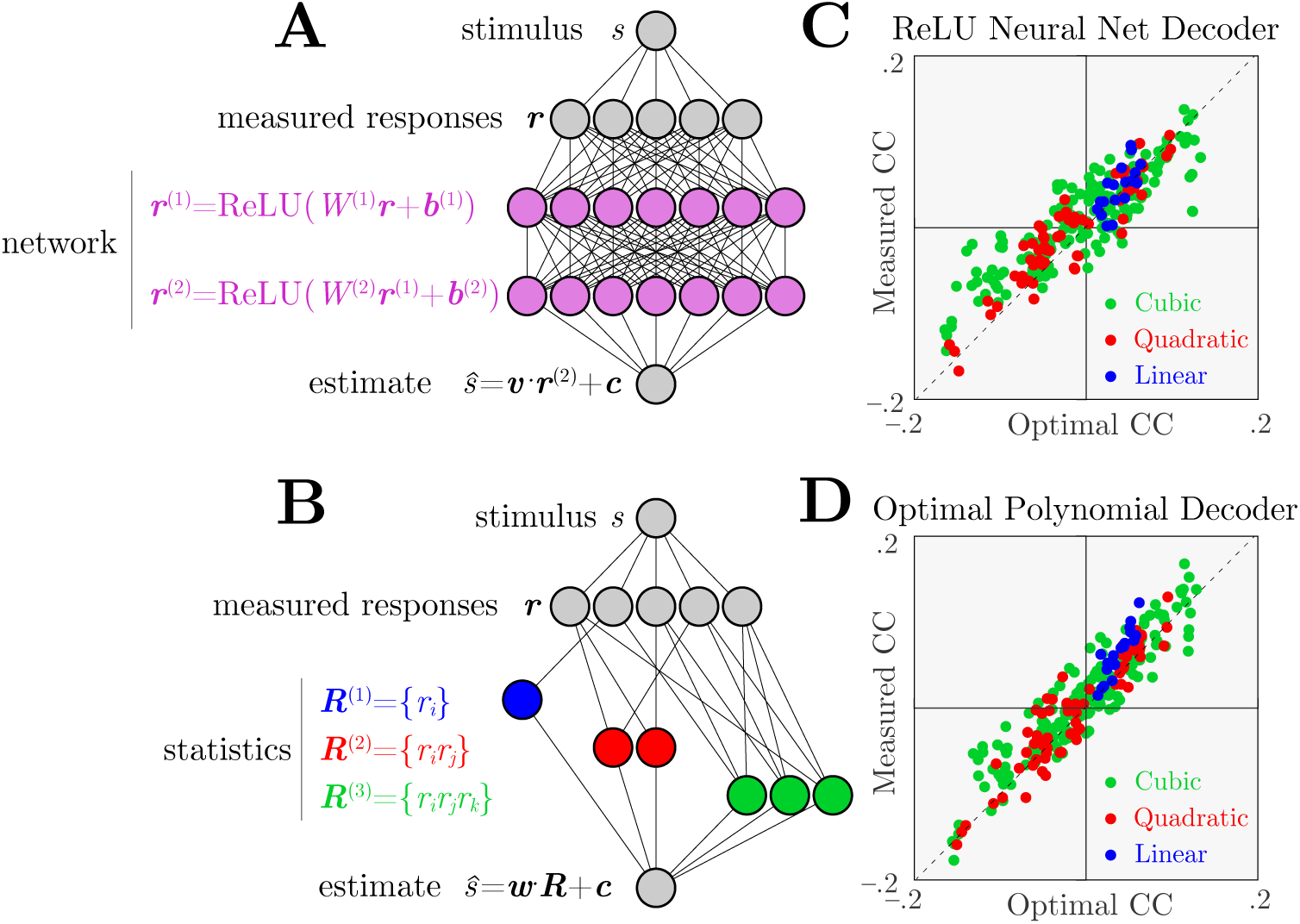
Identifying optimal nonlinear decoding by a generic neural network using nonlinear choice correlations. Neural responses ***r*** are constructed to encode stimulus information in polynomial sufficient statistics up to cubic order (Methods Eq. 13). These responses are decoded by an artificial nonlinear neural network or polynomial nonlinearities, and we evaluate the quality of the decoding using polynomial nonlinearities for both cases. (**A**) Architecture of a network that uses ReLU nonlinearities trained to extract the relevant information. (**B**) Architecture of a second network that instead uses polynomial nonlinearities to extract the relevant information. (**C, D**) Choice correlations based on polynomial statistics show that both networks’ computations are consistent with optimal nonlinear decoding (Methods 4.2.2), even though the simulated networks used different implementations to extract the stimulus information. Horizontal axis shows optimal choice correlations (Eq. 7); vertical axis shows measured choice correlations (Eq. 5).

#### 2.2.4 Redundant codes

It might seem unlikely that the brain uses optimal, or even near-optimal, nonlinear decoding. Even if it does, there are an enormous number of high-order statistics for neural responses, so the information content in any one statistic could be tiny compared to the total information in all of them. For example, with *N* neurons there are on the order of *N*^2^ quadratic statistics, *N*^3^ cubic statistics, and so on. With so many statistics contributing information, the choice correlation for any single one would then be tiny according to the ratio in Eq. 7, and would be indistinguishable from zero with reasonable amounts of data. Past theoretical studies have described nonlinear (specifically, quadratic) codes with extensive information that grows proportionally with the number of neurons [2, 28]. This would indeed imply immeasurably small choice correlations for large, optimally decoded populations.

A resolution to these concerns is information-limiting correlations [3]. The past studies that derive extensive nonlinear information treat large cortical populations in isolation from the smaller sensory population that would naturally provide its input [2, 28]. Yet when a network inherits information from a much smaller input population, the expanded neural code becomes highly redundant: the brain cannot have more information than it receives [50]. Noise in the input is processed by the same pathway as the signal, and this generates noise correlations that can never be averaged away [3].

Previous work [3] characterized linear information-limiting correlations for fine discrimination tasks by decomposing the noise covariance into Σ = Σ_0_ + *ϵ****f***′***f***′^⊺^, where *ϵ* is the variance of the information-limiting component and Σ_0_ is noise that can be averaged away with many neurons.

For *nonlinear* population codes, it is not just the mean that encodes the signal, ***f*** (*s*) = ⟨***r***|*s*⟩, but rather the nonlinear statistics ***F*** (*s*) = ⟨***R***(***r***)|*s*⟩. Likewise, the noise does not comprise only second-order covariance of ***r***, Cov(***r***|*s*), but rather the second-order covariance of the relevant nonlinear statistics, Γ = Cov(***R***(***r***)|*s*) (Section 2.1.3). Analogous to the linear case, these correlations can be locally decomposed as

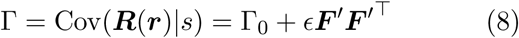

where *ϵ* is again the variance of the information-limiting component, and Γ_0_ is any other covariance which can be averaged away in large populations. The information-limiting noise bounds the estimator variance 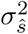 to no smaller than *ϵ* even with optimal decoding.

Neither additional cortical neurons nor additional decoded statistics can improve performance beyond this bound. As a direct consequence, when there are many fewer sensory inputs than cortical neurons, many distinct statistics *R*_*k*_(***r***) will carry redundant information. Under these conditions, many choice correlations 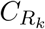 can be substantial even for optimal nonlinear decoding: the discriminabilities 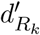 of redundant statistics can be comparable to the discriminability *d*′ of the whole population, producing ratios 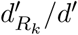 that are not small (Figure 5).

**Figure 5:**
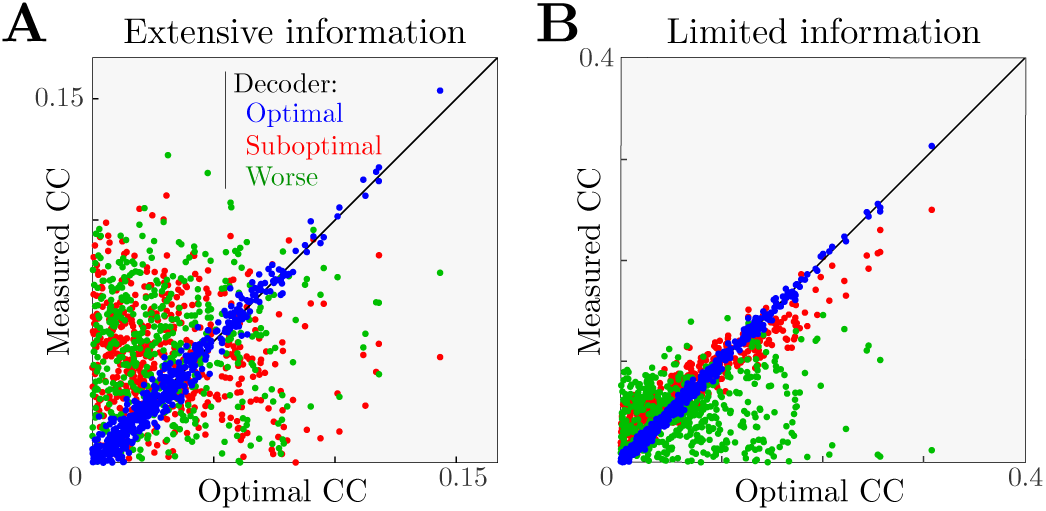
Information-limiting noise makes a network more robust to suboptimal decoding. (**A**) A simulated optimal decoder produces measured choice correlations that match our optimal predictions (blue, on diagonal). In contrast, when a noise covariance Γ_0_ permits the population to have extensive information, then a sub-optimal decoder can exhibits a pattern of choice correlations that does not match the prediction of optimal decoding. Here we show two suboptimal decoders, one that is blind to higher-order correlations (***w*** ∝ ***F***′, red), and another ‘worse’ decoder that has the same weights but with 40% random sign flips (green). As in Figure 4, horizontal axis shows optimal choice correlations (Eq. 7) and vertical axis shows measured choice correlations (Eq. 5). (**B**) When information is limited, the same decoding weights may be less detrimental, and thus exhibit a similar pattern of choice correlations as an optimal decoder (red), or if they are sufficiently bad they may retain a suboptimal pattern of choice correlations (green).

#### 2.2.5 Decoding efficiency revealed by choice correlations

Even if decoding is not strictly optimal, Eq. 7 can be satisfied due to information-limiting correlations. Decoders that seem substantially suboptimal because they fail to avoid the largest noise components in Γ_0_ can be nonetheless dominated by the bound from information-limiting correlations. This will occur whenever the variability from suboptimally decoding the noise Γ_0_ is smaller than the information-limiting variance *ϵ*. Just as we can decompose the nonlinear noise correlations into information-limiting and other parts, we can decompose nonlinear choice correlations into corresponding parts as well, with the result that

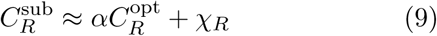

where *χ*_*R*_ depends on the particular type of suboptimal decoding (Supporting Information S.3.2). The slope *α* between choice correlations and those predicted from optimality is given by the fraction of estimator variance explained by information-limiting noise, 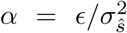. This slope therefore provides an estimate of the efficiency of the brain’s decoding.

Figure 5 shows an example of one decoder that would be suboptimal without redundancy, but is nonetheless close to optimal when information limits are imposed. This rescue of optimality does not happen for all decoders, however. The figure also shows another decoder that is so suboptimal that it throws away most of the available information even when there is substantial redundancy. The patterns of choice correlations reflect this.

In realistically redundant models with more cortical neurons than sensory inputs, many decoders could be near-optimal, as we recently discovered in experimental data for a linear population code [27]. However, even in redundant codes there may be substantial inefficiencies and information loss [51], especially for unnatural tasks [52], so it is scientifically interesting to discover near-optimal nonlinear computation even in a redundant code.

### 2.3 Application to experimental data

#### 2.3.1 Variance discrimination task

We applied our optimality test to data recorded with Utah arrays from primate visual cortex (V1) during a nonlinear decoding task (Method 4.4). Monkeys faced a Two-Alternative Forced Choice task (2AFC) in which they categorized an oriented grating based on whether it came from a wide or narrow distribution of orientations [53] (Figure 6A,B). The categorical target variable *s* is therefore the variance of the orientation distribution.

**Figure 6:**
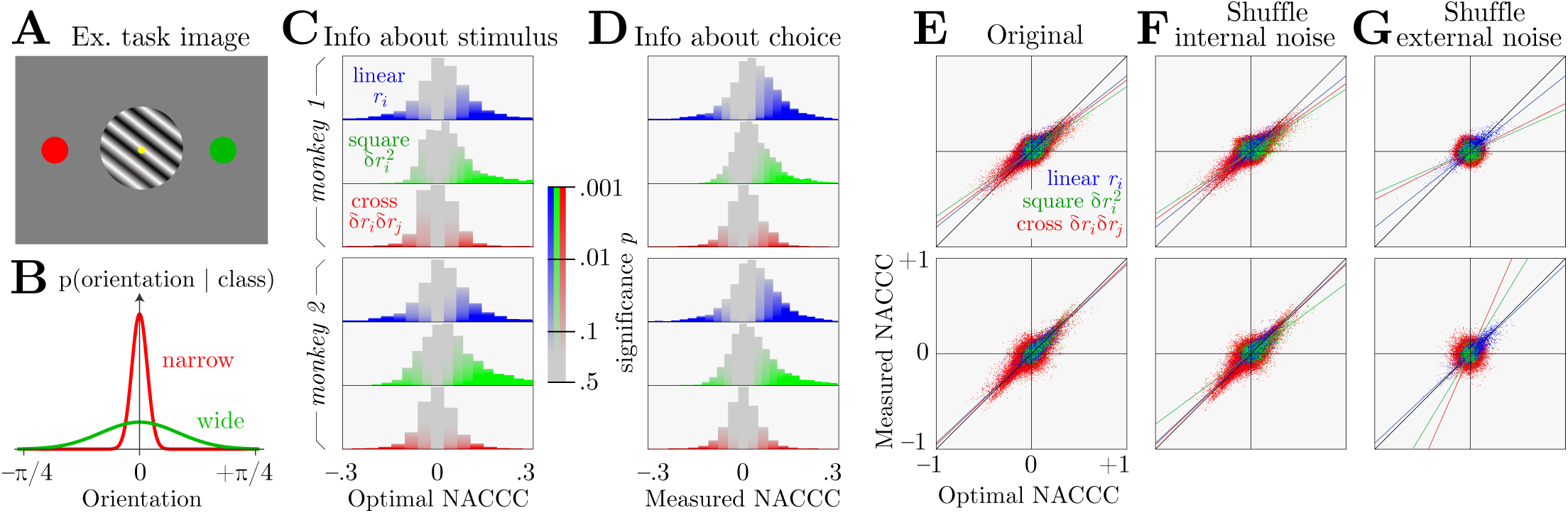
Nonlinear information and choice correlations in a variance discrimination task, for neural data from two monkeys. (**A**) Example oriented grating and saccade targets. (**B**) The orientations of the gratings were drawn from a narrow or wide distribution, and the monkey had to guess which by saccading to the appropriate target. (**C**) Neurons contain linear and nonlinear information about the task variable. This is revealed by the Normalized Average Conditional Choice Correlations (NACCC, Eq. 17) predicted for *optimal* decoding, which are proportional to the measured signal-to-noise ratios (Eq. 7) for each neural response pattern (blue *r*_*i*_, green 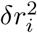, red *δr*_*i*_*δr*_*j*_). Color saturation indicates statistical significance (Methods 4.4). (**D**) These neurons also contain significant information about the animal’s choice, as computed by the *measured* NACCC. (**E**) The measured and optimal NACCCs are highly correlated, with a proportionality near 1 (lines). Each point represents one response pattern (*e.g. δr*_*i*_*δr*_*j*_) in one session. Top and bottom panels are data from two different monkeys. These two plotted quantities are strongly correlated (0.76, 0.65, 0.53 for linear, square and cross terms for monkey 1; 0.80, 0.83, 0.72 for monkey 2). (**F**) Shuffling internal noise correlations while preserving nuisance correlations maintains the relationship between prediction and nonlinear choice correlations, implying that internal noise is not responsible for the correlations. (**G**) Shuffling nuisance correlations across trials (Methods 4.4) nearly eliminates the relationship between measured and predicted nonlinear choice correlations (0.76, 0.05, 0.04 for monkey 1; 0.80, 0.10, 0.11 for monkey 2), implying that nuisance variation creates the nonlinear code.

#### 2.3.2 Evidence for optimal nonlinear computation in macaque brains

This is a coarse discrimination task, since the response statistics change significantly with the stimulus. As for fine discrimination, we again find that when decoding is optimal, random fluctuations in choices are correlated with neural responses to the same degree that those responses can discriminate between stimuli. However, as we describe in the Methods (4.4) and Supplement (S.6.1), this relationship is slightly more complicated for coarse discrimination, since the response statistics change significantly with the stimulus. For this reason we need to use a slightly more complicated measure of choice correlation that we call Normalized Average Conditional Choice Correlation (NACCC, Eq. 17). However, the end result is the same: choice correlations for optimal decoding are equal to the ratio of discriminabilities (Eq. 7; proof in Supplemental Information S.5, Eq. 138). Again there is a correction factor of order 1 for binary choices instead of continuous estimation (Methods 4.2.1, Supplemental Information S.6.4, Eq. 185).

V1 responses contain information about orientation [54]. Here we found that V1 responses also contain some linear information about the orientation *variance* (Figure 6C, blue). Since these neurons have linear information, they have already performed some useful nonlinear transformations of the input within their receptive field.

However, because neural responses in this brain area can be linearly decoded to compute orientation, a good decoder for the orientation variance would naturally be quadratic in those responses. Indeed, we found information in the quadratic statistics of neural responses, 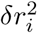 and *δr*_*i*_*δr*_*j*_ (Figure 6C, red and green), suggesting that downstream nonlinear computations could extract additional information from the neural responses. Here we first eliminate the linear information when we compute these neural nonlinear statistics by using *δr*_*i*_ = *r*_*i*_ − ⟨*r*_*i*_|*ŝ*_1_⟩, where *ŝ*_1_ = ***w***_opt_ · ***r*** + *c* is the optimal estimate decoded only from a linear combination of available neural responses.

We also found that these quadratic statistics contained substantial nonlinear information about the behavioral choice (Figure 6D). In general, there is no guarantee that the particular nonlinear statistics that are informative about the stimulus are also informative about the choice. Our theory of optimal decoding predicts specifically that these quantities should be directly proportional to each other.

Indeed, in two monkeys, we found that nonlinear choice correlations were highly correlated with nonlinear information (Figure 6E).

Remarkably, when we compare the measured nonlinear choice correlations to the ratio of discriminabilities after adjusting for the binary data (Methods 4.4), the slopes of this relationship for the two animals were near the value of 1 that Eq. 7 predicts for optimal decoding (Figure 6E).

#### 2.3.3 Controls to find the origins of choice correlations

To evaluate whether internal noise correlations contribute nonlinear information or choice correlations, we created a shuffled data set that removed internal noise correlations while preserving external nuisance correlations. That is, we independently selected responses to trials with matched target stimulus (variance), nuisance (orientation), and choice, and repeated our analysis on these shuffled data (Figure 6F). The observed relationship between predicted and observed choice correlations was the same as in the original test, indicating that nuisance variations were sufficient to drive the nonlinear information and decoding.

We then shuffled the external nuisance correlations by randomly selecting responses to trials with matched target stimulus and choice, but now using *unmatched* nuisance variables, and again repeated the analysis (Figure 6G). In other words, we picked responses from different trials that came from the same signal category (wide or narrow) and elicited the same choice but had different orientations, and we picked these trials (and thus their stimulus orientations) *independently* for neurons *i* and *j*. The strong statistical relationship observed between predicted and measured nonlinear choice correlations vanished with this shuffling, indicating that the nuisance variation was necessary for the nonlinear information and nonlinear decoding.

These shuffle controls removed noise correlations and nuisance correlations, respectively. Combining the conclusions from these controls, we find no evidence that the brain optimally decodes any stimulus-dependent internal noise correlations in this task. Recent analyses of these same data found that internal noise did in fact influence the monkeys’ behavioral choices [55], but this effect was subtle and only apparent when examining the entire neural population simultaneously. In our analysis this effect is buried in the noise, so our method is not sensitive enough to tell if these large-scale patterns induced by internal noise are used optimally or suboptimally. However, we can detect that the brain contains information that is encoded nonlinearly due to external nuisance variation, and that this information is indeed decoded near-optimally by the brain.

One monkey performed slightly worse than an ideal observer, with a probability correct of 0.76, compared to the ideal of 0.82 (Methods 4.4) — even while its decoding was near-optimal, with an efficiency of 0.96±0.04 (mean ± 95% confidence intervals). This suggests that information is lost in the encoding stage somewhere between the stimulus and the recorded neurons, and not downstream of those neurons. The other monkey had similar overall performance (probability correct of 0.74) but worse decoding efficiency (0.75±0.08). This suggests the second monkey’s task performance has limitations arising downstream of the recorded neurons.

## 3 Discussion

This study introduced a theory of nonlinear population codes, grounded in the natural computational task of separating relevant and irrelevant variables. The theory considers both encoding and decoding — how stimuli drive neurons, and how neurons drive behavioral choices. It showed how correlated fluctuations between neural activity and behavioral choices could reveal the efficiency of the brain’s decoding. Unlike previous theories [2, 28], ours remains consistent with biological constraints due to the large cortical expansion of sensory representations by incorporating redundancy through nonlinear information-limiting correlations. Crucially, this theory provides a remarkably simple test to determine if downstream nonlinear computation decodes all that is encoded.

Alternative methods to estimate whether animals decode their information efficiently rely upon comparing behavioral performance to performance of an ideal observer that can access the entire population. Even with impressive advances in neurotechnology, this challenge remains out of reach for large populations. In contrast, our proposed method to test for optimal decoding has a vastly lower experimental burden. It requires only that a few cells be recorded simultaneously while an animal performs a task.

On the negative side, this simple test does not offer a complete description of neural transformations. It instead tests just one important hypothesis about their functional role — that the brain performs optimal decoding. However, the theory does provide a practical way to estimate decoding efficiency. The brain may not be optimal, but instead may be satisfied by a more modest decoding efficiency. In this case, more work is needed to understand which suboptimalities the brain tolerates for satisfactory performance [56].

### 3.1 Which nonlinearities should we test?

If all neural signals are decoded optimally, then all choice correlations for any function of those signals should also be consistent with optimal decoding, since they contain the same information (Figure 4). Yet for the wrong or incomplete nonlinearities that do not disentangle the task-relevant variables from the nuisance variables, the test may be inconclusive, just as it was for linear decoding of a nonlinear code: the chosen nonlinear functions may not extract linearly decodable information nor have any choice correlation.

The optimal nonlinearities would be those that collectively extract the sufficient statistics about the relevant stimulus, which will depend on both the task and the nuisance variables. In complex tasks, like recognizing object from images with many nuisance variables, most of the relevant information lives in higher-order statistics, and therefore require more complex nonlinearities to extract. In such high-dimensional cases, our proposed test is unlikely to be useful. This is because our method expresses stimulus estimates as sums of nonlinear functions, and while that is universal in principle [57], that is not a compact way to express the complex nonlinearities of deep networks. Relatedly, it may be difficult to see statistically significant information or choice correlations for nonlinear statistics that provide many small contributions to the behavioral output. Alternatively, with guidance from trained neural network models, our method could potentially judge whether those nonlinearities provide a good description of neural decoding. This decoding perspective would complement studies that demonstrate a match between the encodings of brains and artificial neural networks [10, 58].

The best condition to apply our optimality test is in tasks of modest complexity but still possessing fundamentally nonlinear structure. Some interesting examples where our test could have practical relevance include motion detection using photoreceptors [59], visual search with distractors (XOR-type tasks) [30, 60], sound localization in early auditory processing before the inferior colliculus [61], or context switching in higher-level cortex [62].

### 3.2 Limitations

For efficient decoding in a learned task, the optimality test (Eq. 7) is necessary but not sufficient. If the brain neglects some of informative sufficient statistics, and we don’t test these neglected statistics either, then we could find the brain is consistent with our optimal decoding test, yet still be suboptimal. Only if the test is passed for *all* statistics will the test be conclusive. For an extreme example, a single neuron might pass the test, but if other neurons don’t, then the brain is not using its information well. On a broader scale, one might find that all individual responses *r*_*k*_ pass the optimality test, while products of responses *r*_*j*_*r*_*k*_ fail. This would be consistent with linear information being used well while distinct quadratic information is present but unused; on the other hand this outcome would not be consistent with quadratic statistics that are uninformative but decoded anyway, since that would increase the output variance beyond that expected from the linear information. Future work will demonstrate how we can use identify properties of suboptimal decoders [56] with nonlinear choice correlations.

Our approach is currently limited to feedforward processing, which unquestionably oversimplifies cortical processing. The approach can be generalized to recurrent networks by considering spatiotemporal statistics [56]. Nonetheless, feedforward models do a fair job of capturing the representational structure of the brain [10].

Feedback could also cause suboptimal networks to exhibit choice correlations that seem to resemble the optimal prediction. If the feedback is noisy and projects into the same direction that encodes the stimulus, such as from a dynamic bias [63], then this could appear as information-limiting correlations, enhancing the match with Eq. 7. This situation could be disambiguated by measuring the internal noise source providing the feedback, though of course this would require more simultaneous measurements.

### 3.3 Choice correlations from internal versus external noise

Since many stimulus-dependent response correlations are induced by external nuisance variation, not internal noise, we might not find informative stimulus-dependent noise correlations upon repeated presentations of a fixed stimulus. Indeed, our analysis found no evidence of internal noise generating nonlinear choice correlations (Figure 6). Those correlations may only be informative about a stimulus in the presence of natural nuisance variation. For example, if a picture of a face is shown repeatedly without changing its pose, then small expression changes can readily be identified by linear operations; if the pose can vary then the stimulus is only reflected in higher-order correlations [9].

In contrast, we *should* see some nonlinear choice correlations even when nuisance variables are fixed. This is because neural circuitry must combine responses nonlinearly to eliminate natural nuisance variation, and any internal noise passing through those same channels will thereby influence the choice. Although they may be smaller and more difficult to detect than the fluctuations caused by the nuisance variation, this influence will manifest as nonlinear choice correlations. In other words, nonlinear noise correlations need not predict a fixed stimulus, but they may predict the choice (Supplementary Information S.4).

For optimal decoding, the choice correlations measured using fixed nuisance variables will differ from Eq. 7, which should strictly hold only when there is natural nuisance variation. This is implicit in Eq. 7, since the relevant quantities are conditioned only on the relevant stimulus *s* while averaging over the nuisance variations *ν* and internal noise. However, under some conditions, a related prediction for nonlinear choice correlations holds even without averaging over nuisance variables (Supplementary Information S.4).

### 3.4 Conclusion

Despite the clear importance of computation that is both nonlinear and distributed, and evidence for nonlinear coding in the cortex [30, 32–34], most neuroscience applications of population coding concepts have assumed linear codes and linear readouts [10, 27, 38, 64, 65]. The few that directly address nonlinear population codes either have an impossibly large amount of encoded information [2, 28], or investigate abstract properties unrelated to structured tasks [66]. Some experimental studies have been able to extract additional information from recorded populations using nonlinear decoders [30, 67], but the inferred properties of such decoders are based on recordings being a representative sample that can be extrapolated to larger populations. Unknown correlations and redundancy prevents that from being a reliable method [23, 68].

Our method to understand nonlinear neural decoding requires neural recordings in a behaving animal. The task must be hard enough that it makes some errors, so that there are behavioral fluctuations to explain. Finally, there should be a modest number of nonlinearly entangled nuisance variables. Unfortunately, many neuroscience experiments are designed without explicit use of nuisance variables. Although this simplifies the analysis, this simplification comes at a great cost, which is that the neural circuits are being engaged far from their natural operating point, and far from their purpose: there is little hope of understanding neural computation without challenging the neural systems with nonlinear tasks for which they are required. In this context, it is especially noteworthy that a mismatch between choice correlations and the optimal pattern might not indicate that the brain is suboptimal, but instead that the nuisance variation in the experimental task may not match the natural tasks the brain has learned. For this reason it is important for neuroscience to use natural tasks, or at least naturalistic ones, when aiming to understand computational function [69].

Our statistical perspective on feedforward nonlinear coding in the presence of nuisance variables provides a useful framework for thinking about neural computation. Furthermore, choice-related activity provides guidance for designing interesting experiments to measure not only how information is encoded in the brain, but how it is decoded to generate behavior.

## Author contributions

XP conceived the theoretical framework. XP and QY designed and performed the mathematical analyses. QY performed the simulations. EW, RJC and AT designed the experiments for a study with Wei Ji Ma; EW, RJC and AT performed the experiments; EW preprocessed the neural data; QY and XP analyzed the neural data. QY and XP wrote the manuscript; all authors discussed the results and commented on the manuscript.

## Acknowledgements

The authors thank Jeff Beck, Valentin Dragoi, Arun Parajuli, Alex Pouget, and Haim Sompolinsky for helpful conversations. This work was supported by NSF CAREER grant 1552868 to XP, by NeuroNex grant 1707400 to XP and AT, and by NSF Grant No. PHY-1748958, NIH Grant No. R25GM067110, and the Gordon and Betty Moore Foundation Grant No. 2919.01.

## Data and code availability

The data that support the findings of this study are available from the corresponding author upon reasonable request. All custom code used for electrophysiology data collection and data processing are made publicly available at github.com/atlab. Experimental data for Figure 6 and code used for analysis and figure generation are available for download from github.com/xaqlab/nonlinear_choice_correlation.

## 4 Online Methods

### 4.1 Encoding models

#### 4.1.1 Orientation estimation with varying spatial phase

Figure 1 illustrates how nuisance variation can eliminate a neuron’s mean tuning to relevant stimulus variables, relegating the neural tuning to higher-order statistics like covariances. In this example, the subject estimates the orientation of a Gabor image, *G*(***x***|*s, ν*), where ***x*** is spatial position in the image, and *s* and *ν* are the orientation and spatial phase (nuisance) of the image, respectively (Supplemental Material S.1.1). The model visual neurons are linear Gabor filters like idealized simple cells in primary visual cortex, corrupted by additive white Gaussian noise. Their responses are thus distributed as ***r*** ∼ *P* (***r***|*s, ν*) = *N* (***r***|***f*** (*s, ν*), *ϵI*), whe re *ϵ* is the noise variance and the mean ***f***(*s, ν*) = ⟨***r***|*s, ν*⟩ = ∑_***r***_ ***r*** *p*(***r***|*s, ν*) is determined by the overlap between the image and the receptive field.

When the spatial phase *ν* is known, the mean neural response contains all the information about orientation *s*. The brain can decode responses linearly to estimate orientation near a reference *s*_0_.

When the spatial phase varies, however, each mean response to a fixed orientation will be combined across different phases: ***f*** (*s*) = ⟨***r***|*s*⟩ = ∑_***r***_ ***r*** *p*(***r***|*s*) = ∫ *dν* ∑_***r***_ ***r*** *p*(***r***|*s, ν*)*p*(*ν*). Since each spatial phase can be paired with another phase *π* radians away that inverts the linear response, the phase-averaged mean is ***f***(*s*) = 0. Thus the brain cannot estimate orientation by decoding these neurons linearly; nonlinear computation is necessary.

The covariance provides one such tuned statistic. We define Cov_*ij*_ (***r***|*s, ν*) as the neural covariance for a fixed input image (noise correlations), and Cov_*ij*_ (***r***|*s*) as the neural covariance when the nuisance varies (nuisance correlations). According to the law of total covariance,

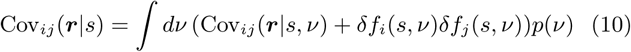

where *δf*_*i*_(*s, ν*) = *f*_*i*_(*s, ν*) − ⟨*f*_*i*_(*s, ν*)⟩_*ν*_. Supplementary Information S.1.1 shows in detail how Cov_*ij*_ (***r***|*s*) is tuned to *s*.

#### 4.1.2 Exponential family distribution and sufficient statistics

It is illuminating to assume the response distribution conditioned on the relevant stimulus (but not on nuisance variables) is approximately a member of the exponential family with nonlinear sufficient statistics,

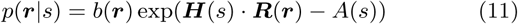

where ***R***(***r***) is a vector of sufficient statistics for the natural parameter ***H***(*s*), *b*(***r***) is the base measure, and *A*(*s*) is the log-partition function. In this case, a finite number of sufficient statistics contains all of the information about the stimulus in the population response, and all other tuned statistics may be derived from them.

Estimation and inference are closely connected in the exponential family. In Supplemental Material S.1.2.2, we show that the optimal local estimation can be achieved by linearly decoding the nonlinear sufficient statistics, *ŝ* = ***w***^⊺^***R***(***r***)+*c*. The decoding weights minimize the variance of an unbiased decoder,

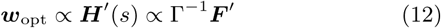

where ***F***′ = *∂* ⟨***R***(***r***)|*s*⟩ */∂s* is the sensitivity of the statistics to changing inputs, and Γ = Cov(***R***|*s*) is the stimulus-conditioned response covariance — which generally includes nuisance correlations (Section 2.1.3).

#### 4.1.3 Quadratic encoding

In a quadratic coding model, the distribution of neural responses is described by the exponential family with up to quadratic sufficient statistics, ***R***(***r***) = {*r*_*i*_, *r*_*i*_*r*_*j*_} for *i, j* ∈ {1, …, *N*}. A familiar example is the Gaussian distribution with stimulus-dependent covariance Σ(*s*). In order to demonstrate the coding properties of a purely nonlinear neural code, here we assume that the mean tuning curve *f*(*s*) is constant, while the stimulus-conditional covariances Σ_*ij*_ (*s*) depend smoothly on the stimulus. We can quantify the information content of the neural population using Equation 61.

#### 4.1.4 Cubic encoding

In our cubic coding model, the distribution of neural responses is described by the exponential family with up to cubic sufficient statistics, ***R***(***r***) = {*r*_*i*_, *r*_*i*_*r*_*j*_, *r*_*i*_*r*_*j*_*r*_*k*_} for *i, j, k* ∈ {1, …, *N*}.

We approximate a three-neuron cubic code first using purely cubic components, and we then apply a stimulus-dependent affine transformation to include linear and quadratic statistics. The pure cubic code is used for a vector ***z*** with sufficient statistics *z*_*i*_*z*_*j*_*z*_*k*_ (and a base measure 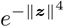 to ensure the distribution is bounded and normalizable).

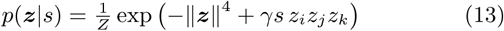

We approximate this distribution by a mixture of four Gaussians. The mixture is chosen to reproduce the tetrahedral symmetry of the cubic distribution (Supplementary Figure S1), which allows the cubic statistics of responses to be stimulus dependent, leaving stimulus-independent quadratic and linear statistics.

To generate larger multivariate cubic codes for Figure (S1), for simplicity we assume the pure cubic terms only couple disjoint triplets of variables, and sample independently from an approximately cubic distribution for each triplet. To convert this purely cubic distribution to a distribution with linear and quadratic information, we shift and scale these cubic samples ***z*** in a manner dependent on *s*:

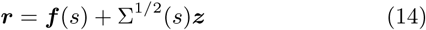

where ***f*** (*s*) and Σ(*s*) describes the desired signal-dependent mean and covariance (see Supplemental Material S.1.4).

### 4.2 Nonlinear choice correlations

For fine discrimination tasks, the nonlinear choice correlation between the stimulus estimate *ŝ* = ***w***^⊺^***R*** + *c* and one nonlinear function *R*_*k*_ (the *k*th element of the vector ***R***) of recorded neural activity ***r*** is

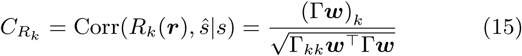

where 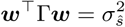 is the estimator variance.

When the relevant response statistics change appreciably over the stimulus range used in the task, such as for the coarse variance discrimination task in Section 2.3), the relevant quantities change slightly. The optimal linear decoder of nonlinear statistics, *ŝ* = ***w*** · ***R*** + *c*, has weights obtained through linear regression:

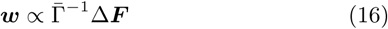

where 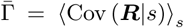 is the average conditional covariance between ***R*** given the stimulus *s*. The differences from Eq 12 are 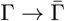 and ***F***′ = *d****F*** */ds* → Δ***F*** */*Δ*s*.

These differences are reflected in a slightly modified measure of correlation that we call Normalized Average Conditional Choice Correlations (NACCC),

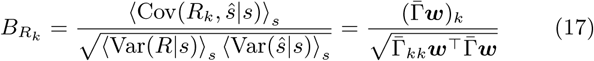

Note that Eq 17 is not actually a correlation coefficient, and may exceed the interval [−1, 1] if Var(*ŝ*|*s*) and Var(*R*_*k*_|*s*) are anticorrelated over *s*. As the stimulus range in a coarse task decreases, and the noise distribution *p*(***R***|*s*) becomes independent of the stimulus, then Eq 17 converges toward Eq 15.

The choice correlation for binary choices differs slightly from that for continuous estimation, for both fine and coarse discrimination tasks, by a factor *ζ* that is typically of order 1 (Supplemental Materials S.6.1).

#### 4.2.1 Optimality test

Substituting the optimal weights (Eq 12) into (15), the optimal nonlinear choice correlation becomes

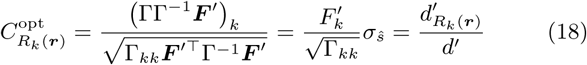

where 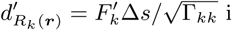 is the fine discriminability provided by *R*_*k*_(***r***) for a stimulus difference of Δ*s*. The same argument holds for coarse discrimination, where 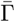 in Eq 17 is canceled by 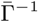 in the optimal weights (Eq 16), yielding 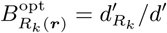.

For fine-scale discrimination, optimal choice correlations can be written in many equivalent ways that reflect the simple relationships between four quantities often used to represent information: discriminability *d*-prime is pro<u>p</u>ortional to the square root of the Fisher information 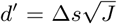 [70]; estimator variance is bounded by the inverse of the Fisher information, 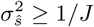; discrimination threshold is proportional to the estimator standard deviation, 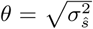 with proportionality given by the threshold condition.

In different experiments (binary discrimination, continuous estimation), it can be most natural to express this optimal decoding prediction as ratios of different measured quantities:

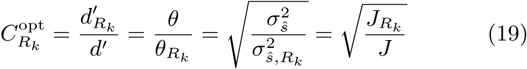

These quantities reflect information between the stimulus and the neural or behavioral responses. Supplemental material S.5 shows how this can be computed easily for general binary discrimination using the total correlation between the responses and the stimuli, 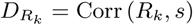, or a continuously varying behavioral choice *ŝ* and the stimuli, *D*_*ŝ*_ = Corr (*ŝ, s*):

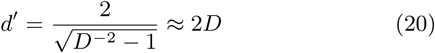

and likewise for 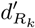. When the behavioral choice is binary rather than continuous, the correlations are modified by a factor *δ* near 1 (Supplemental Information S.6.3, Eq 182). For our experimental conditions, *δ* ≈ 1.2 ± 0.2.

#### 4.2.2 Nonlinear choice correlation to analyze an unknown nonlinearity

In Figure 4, we generated neural responses given sufficient statistics that are polynomials up to third order, ***R***(***r***) = {*r*_*i*_, *r*_*i*_*r*_*j*_, *r*_*i*_*r*_*j*_*r*_*k*_} (Methods 4.1.4). Our model brain decodes the stimulus using a cascade of linear-nonlinear transformations, with Rectified Linear Units (ReLU(*x*) = max(0, *x*)) for the nonlinear activation functions. We used a fully-connected ReLU network with two hidden layers and 30 units per hidden layer. We trained the network weights and biases with backpropagation to estimate stimuli near a reference *s*_0_ based on 20000 training pairs (***r***, *s*) generated by the cubic encoding model. This trained neural network extracted 91% of the information available to an optimal decoder.

### 4.3 Information-limiting correlations

Only specific correlated fluctuations limit the information content of large neural populations [3]. These fluctuations can ultimately be referred back to the stimulus as ***r*** ∼ *p*(***r***|*s* + *ds*), where *ds* is zero mean noise, whose variance 1*/J*_∞_ determines the asymptotic variance of any stimulus estimator. These information-limiting correlations for nonlinear computation can be characterized by the covariance of the sufficient statistics, Γ = Cov(***R***|*s*) conditioned on *s*; the information-limiting component arises specifically from the signal covariance Cov(***F*** (*s*)|*s*). Since the signal for local estimation of stimuli near a reference *s*_0_ is 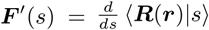, the information-limiting component of the covariance is proportional to ***F***′***F***′^⊺^:

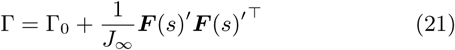

Here Γ_0_ is any covariance of ***R*** that does *not* limit information in large populations. Substituting this expression into (61) for the nonlinear Fisher Information, we obtain

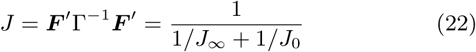

where 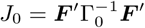 is the nonlinear Fisher Information allowed by Γ_0_. When the population size grows, the extensive information term *J*_0_ grows proportionally, so the output information will asymptote to *J*_∞_.

### 4.4 Application to neural data

All behavioral and electrophysiological data were obtained from two healthy, male rhesus macaque (*Macaca mulatta*) monkeys (L and T) aged 10 and 7 years and weighting 9.5 and 15.1 kg, respectively. All experimental procedures complied with guidelines of the NIH and were approved by the Baylor College of Medicine Institutional Animal Care and Use Committee (permit number: AN-4367). Animals were housed individually in a room located adjacent to the training facility on a 12h light/dark cycle, along with around ten other monkeys permitting rich visual, olfactory, and auditory social interactions. Regular veterinary care and monitoring, balanced nutrition and environmental enrichment were provided by the Center for Comparative Medicine of Baylor College of Medicine. Surgical procedures on monkeys were conducted under general anesthesia following standard aseptic techniques.

Monkeys faced a Two-Alternative Forced Choice (2AFC) to guess whether an oriented drifting grating stimulus came from a narrow or wide distribution of orientations, centered on zero with standard deviations *σ*_+_ = 15° and *σ*_−_ = 3°. Visual contrast was set to 64%. Each trial was initiated by a beeping sound and the appearance of a fixation target (0.15° visual angle) in the center of the screen. The monkey fixated on a fixation target for 300ms within 0.5°–1° visual angle. The stimulus appeared at the center of the screen. After 500ms, colored targets appeared randomly on the left and right, and the monkey then saccades to one of these targets to indicate its choice (red and green targets correspond to narrow and wide distributions).

After the monkey was fully trained, we implanted a 96-electrode microelectrode array (Utah array, Blackrock Microsystems, Salt Lake City, UT, USA) with a shaft length of 1 mm over parafoveal area V1 on the right hemisphere. The neural signals were pre-amplified at the head stage by unity gain preamplifiers (HS-27, Neuralynx, Bozeman MT, USA). These signals were then digitized by 24-bit analog data acquisition cards with 30 dB onboard gain (PXI-4498, National Instruments, Austin, TX) and sampled at 32 kHz. The spike detection was performed offline according to a previously described method [12, 71]. Code for spike detection is available online at github.com/atlab/spikedetection. For each behavioral session and in both monkeys, 95 multiunit neural responses *r*_*k*_ were measured by spike counts in the 500 ms preceding the saccade target onset.

The animals did not perform well on all days, so for further analysis we selected sessions where the performance exceeded 0.7 for monkey 1 (85% of all sessions) and 0.75 for monkey 2 (68% of all sessions).

The task-relevant stimulus *s* is the large or small variance 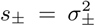 of the distribution over orientations. Orientation is a nuisance variable *ν*, drawn from *p*(*ν*|*s*) = 𝒩(*ν*|0, *s*), which has sufficient statistics that are quadratic in *ν*. If the orientation itself can be estimated locally from linear functions of the neural responses, then the stimulus can be decoded quadratically from those neural responses, 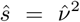. A binary guess about the variance is given by 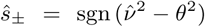 where *θ* is the animal’s orientation threshold. This threshold is optimal where the two stimuli are equally probable: *p*(*ν*|*s*_+_) = *p*(*ν*|*s*_−_), implying that 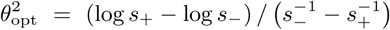. The probability of correctly guessing the orientation variance is 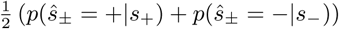, where these probabilities can be computed from the cumulative normal distribution on the correct side of the optimal orientation threshold, 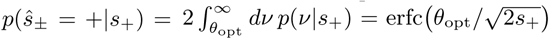; similarly, 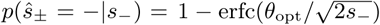. Using values of *s*_±_ for our task, this gives an optimal fraction correct of 0.82.

We computed choice correlations using NACCC (Eq 17), and discriminability based on total correlations between stimulus and response (Eq 20). We adjusted the optimal prediction by constant factors *ζ* and *δ* to account for binary choices using the equations in Supplement S.6.4, with thresholds estimated by logistic regression between choice and the absolute value of the stimulus orientation. We estimated the slopes of the relationship between measured and predicted choice correlation using the angle of the principal component of the bivariate data. We computed standard deviations for these quantities by bootstrapping 100 times.

For our two shuffle controls testing whether correlations between neurons were informative about the stimulus or choice, we selected responses independently from *r*_*i*_ ∼ *p*(*r*_*i*_|*s, ν, ŝ*) (Figure 6F) or *r*_*i*_ ∼ *p*(*r*_*i*_|*s, ŝ*) (Figure 6G).

We evaluate statistical significance of the measured and predicted optimal choice correlations using *p*-values for null distributions based on 100 shuffled choices and 100 shuffled stimuli, while preserving correlations between neural responses. Both null distributions are approximately Gaussian with zero means, so we compute the *p*-value of the choice correlations with respect to the corresponding Gaussian, 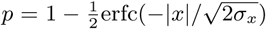 where *x* is the quantity of interest and *σ*_*x*_ is its standard deviation (Figure 6C,D).

## Supplemental material

### S.0 Overview

This supplemental material contains mathematical details and proofs of the central ideas presented in the main text.

**S.1** Encoding models
  **S.1.1** Orientation estimation task with phase as nuisance
  **S.1.2** Exponential family distributions
  **S.1.3** Quadratic codes
  **S.1.4** Cubic codes
**S.2** Information-limiting correlations
**S.3** Analyzing decoding quality
**S.4** Choice correlations from internal and external sources
**S.5** Coarse discrimination and choice correlations
**S.6** Orientation variance discrimination task

### S.1 Encoding models

#### S.1.1 Orientation estimation task with varying spatial phase

In Figure 2B, the subject’s task is to estimate orientation *s* near a reference *s*_0_, based on images *G* of Gabor patterns given by

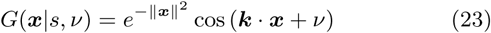

where ***k*** = *κ*(cos *s*, sin *s*). Here the target *s* is the orientation of the pattern, *ν* is a nuisance variable reflecting the spatial phase, ***x*** is the pixel location in the image, and ***k*** is a spatial frequency vector with amplitude *κ* = ‖***k***‖. We assume the spatial receptive field of simple cell *j* in primary visual cortex is also described by a Gabor function

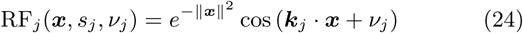

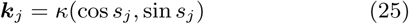

where each neuron has a preferred orientation *s*_*j*_, spatial phase *ν*_*j*_, and spatial frequency ***k***_*j*_. Here for simplicity we assume that all neurons’ preferred spatial frequencies have the same amplitude *κ* that matches the input image.

We model the mean neuronal responses by the overlap between the image and their linear receptive field. This overlap determines the tuning curve of each neuron:

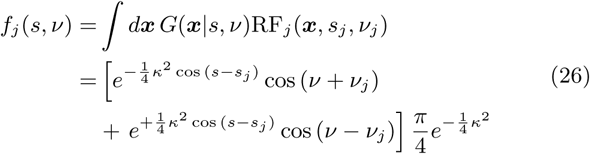

This expression can be written in the form:

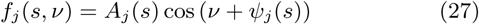

using the stimulus-dependent response amplitude

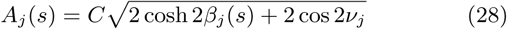

and phase

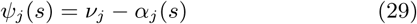

where we define the quantities

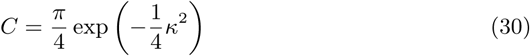

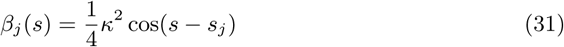

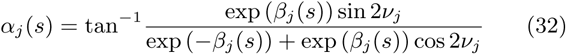

Equation 27 reveals that the mean response of each neuron traces out a sinusoidal oscillation in *ν*, where the amplitude and phase depend on *s* and the specific neuron *j*. The mean tuning for each pair of neurons therefore traces out an ellipse as a function of the nuisance variable, the input’s spatial phase. When we *average* over the ellipse generated by the nuisance variable *ν*, the mean tuning to *s* is abolished — but the response *covariances* (nuisance correlations) remain tuned to *s*.

Assuming each neuron’s response variability is drawn independently from a standard Gaussian 𝒩 (0, 1), we can write the response distribution as

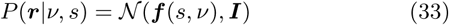

If the spatial phase *ν* were fixed and known, the brain could estimate the orientation just from the mean tuning of the neural responses. However, if the spatial phase is unknown and varies between stimulus presentations uniformly from 0 to 2*π*, the mean tuning ***f*** (*s*) can be expressed as

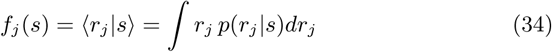

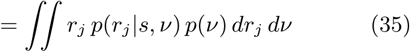

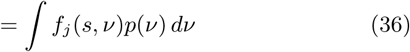

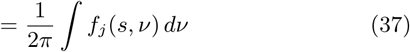

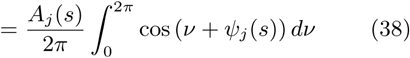

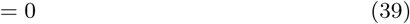

This shows that there is no signal in the mean responses.

However, the brain can perform quadratic computations to eliminate the nuisance variable. We can define Cov_*ij*_ [***r***|*s, ν*] as the neural covariance (noise correlations) when everything in the image is fixed, and Cov_*ij*_ [***r***|*s*] as the neural covariance when the nuisance is unknown and free to vary (nuisance correlations).

Then Cov_*ij*_ [***r***|*s*] is

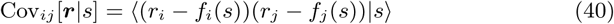

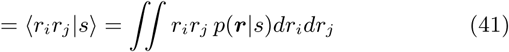

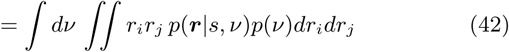

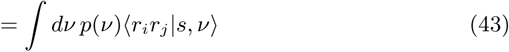

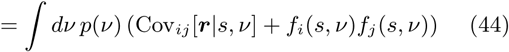

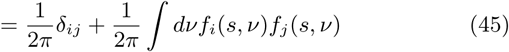

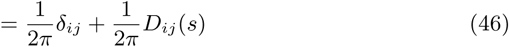

where *D*_*ij*_ (*s*) is given by

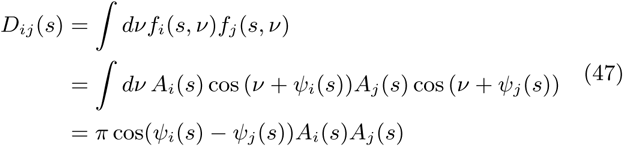

Here when we compute Equation 47, we used the trigonometric identity: 2 cos(*x*) cos(*y*) = cos(*x* + *y*)+ cos(*x* − *y*), and ∫ cos(2*ν* + *ψ*_*i*_ + *ψ*_*j*_)*dν* = 0.

This demonstrates that the neural covariance Cov_*ij*_ [***r***|*s*] depends on the orientation *s*. While linear computation is useless for estimating orientation since the mean responses are untuned (34), quadratic (or higher-order) nonlinear computations can be used to estimate the orientation.

#### S.1.2 Exponential family distributions

For a stimulus *s* and a response ***r***, the conditional probability is a member of the exponential family when

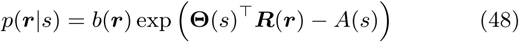

where **Θ**(*s*) are the natural parameters, ***R***(***r***) are the sufficient statistics, *A*(*s*) and *b*(***r***) are the log normalizer and base measure. The statistics ***R***(***r***) are called sufficient because they contain all the information needed to estimate the stimulus *s*.

##### S.1.2.1 Fisher information

One measure of information content that a population response contains about a stimulus is the Fisher information *J* (*s*) [1–3, 24–26]. The Fisher information is given by

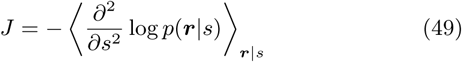

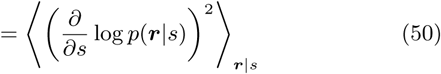

For distributions *p*(***r***|*s*) in the exponential family with sufficient statistics ***R***(***r***), we can compute these quantities analytically. We denote the mean of the sufficient statistics as ***F*** (*s*) = ⟨***R***(***r***)|*s*⟩. This mean ⟨***R***|*s*⟩ can be obtained by differentiating *A*(*s*) by the natural parameters **Θ**(*s*),

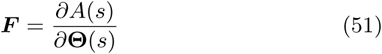

Equation 51 can give us the first and second derivatives of *A*(*s*) over *s*.

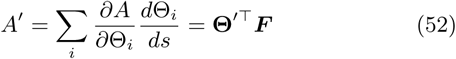

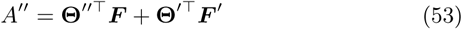

Thus we can compute two definitions of Fisher information.

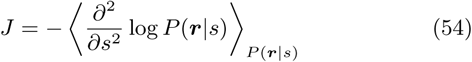

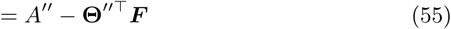

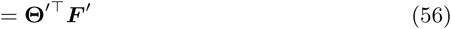

and

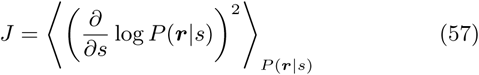

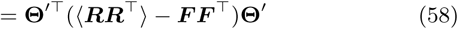

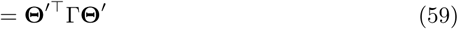

where Γ = Cov[***R***(***r***)|*s*].

Since the two definition are equivalent, we have

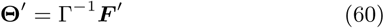

Substituting Equation 60 into Equation 59, we find the Fisher Information for the exponential family [23]

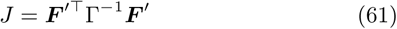

##### S.1.2.2 Optimal estimation in the exponential family

Again assuming responses come from this distribution, we want to compute the maximum likelihood stimulus, *ŝ*, near a reference stimulus *s*_0_:

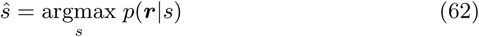

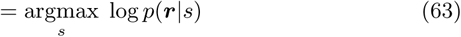

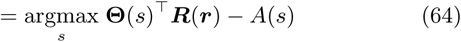

A Taylor expansion around the reference yields

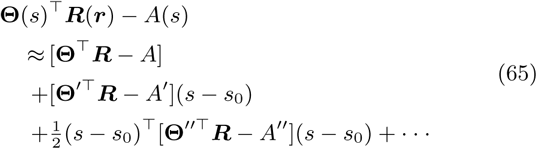

where all functions and derivatives are evaluated at *s*_0_. We find the maximum *ŝ* by differentiating with respect to *s* and setting the result equal to zero:

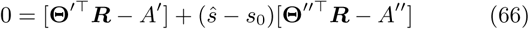

The solution is

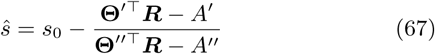

Since ***r*** is a random quantity, we can express ***R*** as a mean and a deviation away from that mean: ***R*** = ⟨***R***|*s*_0_⟩ + *δ****R*** = ***F*** + *δ****R***. In this case, **Θ**″^⊺^***R*** − *A*″ = **Θ**″^⊺^***F*** − *A*″ + **Θ**″^⊺^*δ****R***, where the mean term is precisely the negative Fisher Information −*J* (*s*_0_). If the trial-to-trial fluctuations in the uncertainty are small relative to the average uncertainty then this Fisher term will dominate.

Then we have

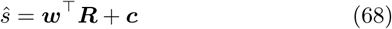

where

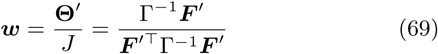

and where we used the results from Equations 61 and 60, with Γ = Cov(***R***|*s*_0_) and ***F*** = ⟨***R***|*s*_0_⟩. Thus, in this limit, the optimal estimator for *s* is a linear decoding of the sufficient statistics ***R***(***r***).

#### S.1.3 Quadratic codes

In a purely quadratic coding model (no linear information), the distribution of neural responses is described by the exponential family with quadratic sufficient statistics, *p*(***r***|*s*) ∼ exp [**Θ**(*s*)^⊺^ ***R***(***r***)] where ***R***(***r***) = (…, *r*_*i*_*r*_*j*_, …). A familiar example is a Gaussian distribution with stimulus-dependent covariance: *p*(***r***|*s*) = *N* (***f***, Σ(*s*)).

As a concrete example we construct a covariance that rotates with stimulus *s*. Any covariance matrix needs to be positive semidefinite. We build Σ(*s*) by setting the eigenvalues to be positive and *s*-independent and eigenvectors to form an orthogonal basis that rotates with *s*:

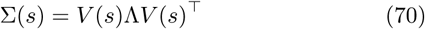

where *V*(*s*) = exp *As* is a rotation matrix in which *A* = −*A*^⊺^ is a real antisymmetric matrix with pure imaginary eigenvalues, and Λ is a diagonal matrix composed of all positive eigenvalues of Σ(*s*).

To calculate the Fisher Information (Equation 61), we need to first calculate the derivative of the mean 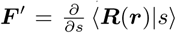 and covariance Γ = Cov[***R***(***r***)|*s*] of the quadratic sufficient statistics.

Because the mean of ***r*** is not dependent on the stimulus in this example, we can compute 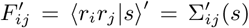, where 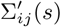 is the derivative of the covariance of ***r***,

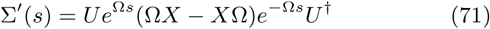

where † denotes a conjugate transpose. Here Ω is a diagonal matrix of eigenvalues for *A, U* is an orthogonal matrix of the eigenvectors of *A*, and *X* = *U*^†^Λ*U*.

The elements in Γ can be expressed as Γ_*ij,kn*_ = ⟨*r*_*i*_*r*_*j*_*r*_*k*_*r*_*n*_|*s*⟩ − ⟨*r*_*i*_*r*_*j*_|*s*⟩ ⟨*r*_*k*_*r*_*n*_|*s*⟩. We can use the following identity for a Gaussian to compute this fourth-order quantity:

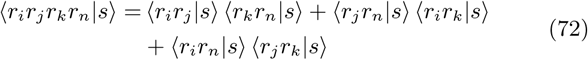

where

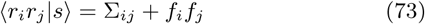

Substitution of the response covariance (Equation 70) into Equation 72 allows us to calculate the covariance Γ of the quadratic sufficient statistics, and thereby to estimate the stimulus and Fisher information for this quadratic code.

#### S.1.4 Cubic codes

In Figure S1 we assume the brain encodes the stimulus using a cubic code. A simple cubic code in ***z*** = (*z*_*i*_, *z*_*j*_, *z*_*k*_) ∈ ℝ^3^ can be written as

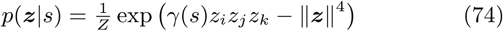

where we include the base measure 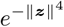 to ensure normalizability (Figure S1A).

For mathematical convenience, we approximate this code by a mixture of Gaussians.

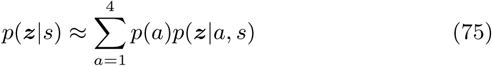

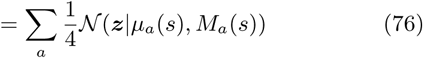

where

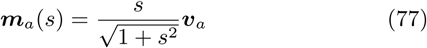

and

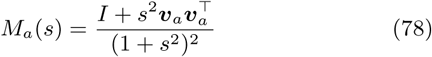

The vectors ***v***_*a*_ reflect the four corners of the tetrahedron, *v*_*a,i*_ = ±1, to match the tetrahedral symmetry of the pure cubic code (Equation 74, Figure S1). To sample from this distribution, we randomly choose a component *a* and then sample from the gaussian 𝒩 (***z***|***m***_*a*_(*s*), *M*_*a*_(*s*)) conditioned on that component.

This distribution has zero mean and identity covariance but a nontrivial skewness tensor, and qualitatively matches the corresponding distribution for the true exponential family distribution with cubic sufficient statistics (Figure S1).

**Figure S1:**
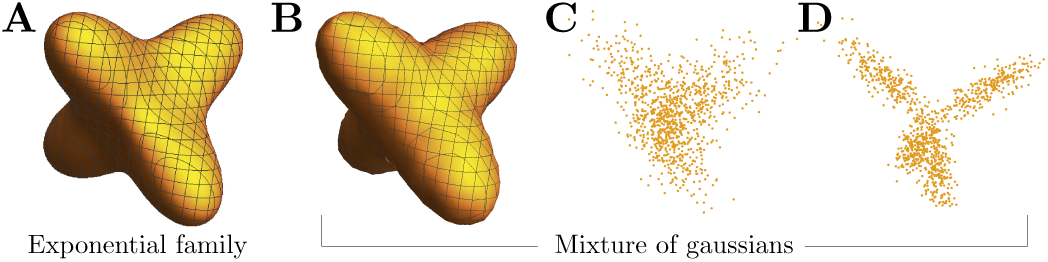
Multivariate skewed distributions. (**A**) Isoprobability contour of an exponential family distribution with cubic statistics in three dimensions, drawn from *p*(***z***|*s*) ∝ exp (*sz*_1_*z*_2_*z*_3_ − ‖***z***‖^4^). (**B**) Isoprobability contour for a mixture of four gaussians (Is 76). (**C**,**D**) Samples drawn from the mixture form, with *s* = 1, 2.

For simplicity, we consider pure cubic codes with non-overlapping cliques of three variables.

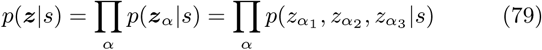

To convert this purely cubic distribution into a distribution with linear and quadratic information as well, we simply shift and scale the distribution in a manner dependent on *s*:

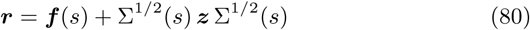

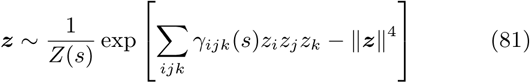

These affine transformations can be incorporated directly into each component of the mixture of gaussians,

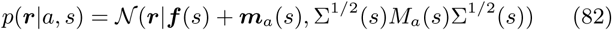

Note that the linear and quadratic information terms are independent of the component *a*.

### S.2 Information-limiting correlations

Information-limiting correlations [3] describe variability that cannot be averaged away because they are indistinguishable from changes in the stimulus. These fluctuations can ultimately be referred back to the stimulus, to appear as ***r*** ∼ *p*(***r***|*s* + *ds*), where *ds* is zero mean noise with variance 1*/J*_∞_ which determines the uncertainty of stimulus. Applying the law of total covariance, we can decompose the covariance of nonlinear statistics ***R***(***r***) conditioned on the stimulus into two parts:

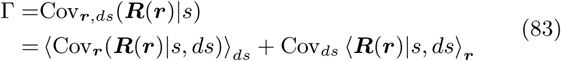

where ⟨·⟩ indicates an expectation value over the subscripted variable. The first term can be computed as follows,

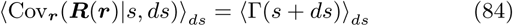

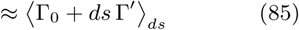

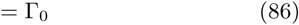

Here we denote the covariance of ***R***(***r***) given *s* and *ds* as Γ(*s*+*ds*). The second equality used a Taylor expansion of Γ(*s* + *ds*) around *s*. The third equality used the fact that the mean of *ds* is zero. Γ_0_ is the covariance of ***R*** in the absence of information-limiting correlations. The second term in Equation 83 can be expressed as

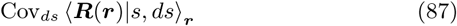

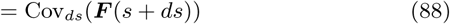

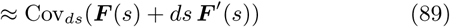

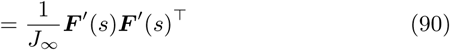

Here we have written the mean of ***R***(***r***) given *s* and *ds* as ***F*** (*s* + *ds*). The second equality used a first-order expansion of ***F*** (*s*+*ds*) around *s*. The third equality used the fact that the variance of *ds* is 1*/J*_∞_.

Equation 83 can therefore be written as

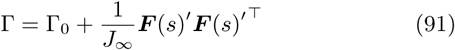

which is a rank-one perturbation of the covariance Γ_0_.

To compute the nonlinear Fisher Information, *J*_*R*(***r***)_ = ***F***′^⊺^Γ^−1^***F***′, we can use the Sherman-Morrison lemma to compute Γ^−1^:

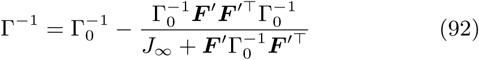

Substituting these equations into the nonlinear Fisher Information (Equation 61) and simplifying, we obtain

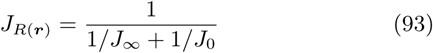

Here 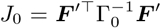 is the nonlinear Fisher Information in the absence of information-limiting correlations. When the population size grows, the term *J*_0_ grows proportionally [1, 2], so for large populations the output information saturates at *J*_∞_.

### S.3 Analyzing decoding quality

#### S.3.1 Unknown nonlinearities

The true nonlinearity that the brain uses to estimate the stimulus is unknown. Thus a crucial question in our decoding analysis is, which nonlinearities to consider? One reasonable set is polynomials in ***r***, *i.e.* a Taylor series expansion of the neural nonlinearities, **Ψ**(***r***) = (*r*_*i*_, *r*_*i*_*r*_*j*_, *r*_*i*_*r*_*j*_ *r*_*k*_, …).

The locally optimal decoder is a weighted sum of the sufficient statistics ***R***(***r***) (Equation 68):

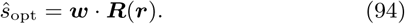

However, the brain might choose a different nonlinear basis ***g***(***r***):

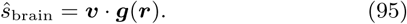

As long as the brain’s nonlinear function spans the same function basis as the sufficient statistics, we can still get all of the information about stimulus from neural population. This allows us to use choice correlation between brain’s estimate *ŝ*_brain_ and our analysis nonlinearity Ψ(***r***) to check the optimality condition (Equation 7).

In Figure 4, we assumed that the optimal nonlinear basis function ***R*** is polynomial nonlinearity up to third order, ***R***(***r***) = (*r*_*i*_, *r*_*i*_*r*_*j*_, *r*_*i*_*r*_*j*_ *r*_*k*_, …). We used cubic codes described in Methods 4.1.4 to generate neural responses for which ***R***(***r***) are sufficient statistics for the stimulus. In this simulation, 18 neuronal responses (six cliques of size 3) were generated using cubic codes.

Our model brain decodes the stimulus using a cascade of linear-nonlinear transformations, with Rectified Linear Units (ReLU(*x*) = max(0, *x*)) for the nonlinear activation functions. We used a fully-connected ReLU network with two hidden layers and 30 units per hidden layer,

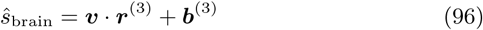

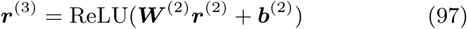

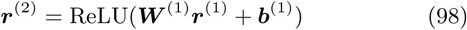

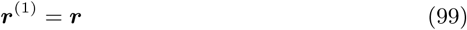

We trained the neural network with 20000 response samples generated from a cubic code driven by stimuli near the reference *s*_0_. We optimized the estimation performance for the neural network using backpropagation to find weights {***W***^(*ℓ*)^}, biases {***b***^(*ℓ*)^}, and readout vector ***v*** that minimized the mean squared error. Our trained neural network performed near-optimally, extracting 91% of the Fisher information compared to optimal decoding based on the true sufficient statistics.

Feigning ignorance of our simulated brain’s true decoder, we applied the nonlinear choice correlation test (Equation 7) using monomial nonlinearities Ψ(***r***) up to third order, *e.g. r*_*i*_, *r*_*i*_*r*_*j*_, 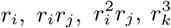, etc. The simulated choice correlations were calculated by Equation 5, where ***R***(***r***) = Ψ(***r***) based on neural responses driven by the reference stimulus *s*_0_, and the stimulus estimate was *ŝ*_brain_. The optimal choice correlation is computed using Equation 7, where 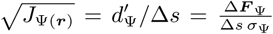, and 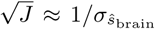. We computed Δ***F***_Ψ_ based on neural population responses ***r***_+_and ***r***_−_ driven by stimuli *s*_+_ = *s*_0_ ± Δ*s/*2. The change in mean was Δ***F***_Ψ_ = ⟨Ψ(***r***_+_)⟩ − ⟨Ψ(***r***_−_)⟩, and the average variance was 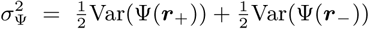. The trained neural network’s estimate *ŝ*_brain_ has a variance 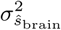 near the reference stimulus *s*_0_. Based on these quantities, Figure 4 shows that we can successfully identify that the brain is near-optimal.

#### S.3.2 Decoding efficiency

A decoder that would be suboptimal for one population code could be near-optimal in the presence of information-limiting noise. In this case, nonlinear choice correlations can be decomposed into a sum of two terms, one from the information-limiting component and the other from the rest of the noise [27]:

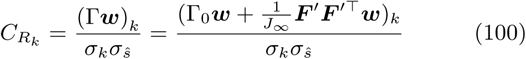

For unbiased decoding, ***w***^⊺^***F***′ = 1. Some manipulation gives [27]

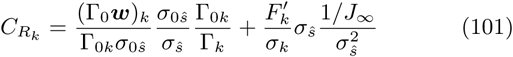

where Γ_0*k*_ = (Γ_0_)_*kk*_ ≈ Γ_*kk*_ for small information-limiting noise variance 1*/J*_∞_ ≪ Γ_0*k*_ (which nonetheless can have a large effect on information despite the small variance), and where *σ*_0*ŝ*_ is the standard deviation of the estimate produced by the same suboptimal decoder ***w*** in the absence of information-limiting correlations, *i.e.* when the covariance of the sufficient statistics is Γ_0_. The variance of *ŝ* can itself be decomposed into two terms as well:

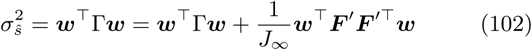

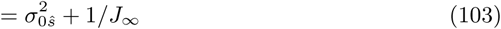

where we assume unbiased decoding, which implies ***w***^⊺^***F***′ = 1. This expression allows us to represent the ratio 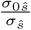 as

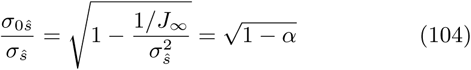

with 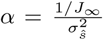. Substituting these into (Eq 101) we find that the choice correlation for a suboptimal decoder in the presence of information-limiting correlations is a weighted sum of the choice correlations for optimal and suboptimal decoding:

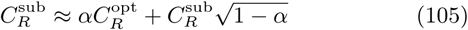

Here 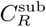 and 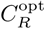 are, respectively, the choice correlations for suboptimal decoding without information-limiting noise (so Γ = Γ_0_), and choice correlations for optimal decoding.

The slope *α* between choice correlations and those predicted from optimal decoding is equal to the fraction of estimator variance explained by information-limiting noise. This slope therefore provides an estimate of the efficiency of the brain’s decoding.

### S.4 Choice correlations from internal versus external noise

The response covariance that drives fluctuations in choices could arise from internal or external (nuisance) variability, or both. Choice correlations predicted for optimal decoding differ depending on whether we condition on the nuisance variables or not. In the main text, we described optimal choice correlations under the distribution *p*(***r***|*s*). This includes variations caused by external nuisance variables, which is sensible since this is what the brain’s decoder must handle. However, it is also potentially informative to examine how purely internal variability correlates with choice, as this is often how choice correlations are assessed. In this section, we derive the choice correlations driven by purely internal noise, for a decoder that learned to remove external nuisance variation as well.

For simplicity we assume that the nonlinear sufficient statistics ***R***(***r***) are linearly tuned to both the stimulus *s* and a scalar nuisance variable *ν*,

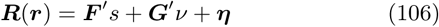

where ***F***′ and ***G***′ characterize the sensitivity of ***R***(***r***) to stimulus *s* and nuisance *ν*, and an internal noise source ***η*** has zero mean with covariance *H*. We assume the brain has a prior over the nuisance variation, *p*(*ν*), with zero mean and variance *ξ*. The total covariance for internal and external fluctuations is then

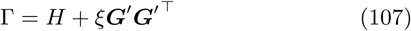

When we measure choice correlations while fixing the nuisance variables in the experiment, we assume the brain retains its decoding strategy accounting for both internal noise and unknown nuisance variation, and not the optimal decoding strategy when the nuisance is fixed and known. These decoding weights are

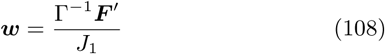

where the denominator *J*_1_ = ***F***′^⊺^Γ^−1^***F***′ is the Fisher information about *s* when there is natural nuisance variation following *p*(*ν*). For distributions in the exponential family, this information saturates the Cramer-Rao bound on an estimator’s variance, so that 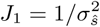. [72] The normalization by *J*_1_ ensures the decoding is locally unbiased. These weights are used to estimate the stimulus according to

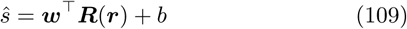

Choice correlations in this fixed-nuisance experiment will be denoted by a lowercase *c*:

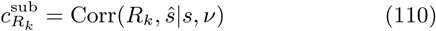

We include the superscript *c*^*sub*^ as a reminder that these choice correlations do not follow the optimal pattern when the decoder is not matched to only the purely internal variability, as here.

We can express these choice correlations as:

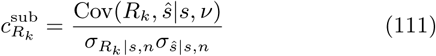

The covariance between *ŝ* and ***R*** is

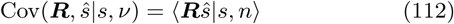

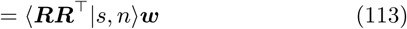

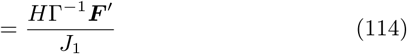

For the scalar nuisance variable we assume here, we can use the Sherman-Morrison lemma to decompose the inverse of the total covariance into a rank-one perturbation of the internal noise inverse covariance:

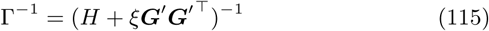

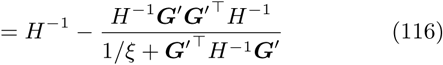

Substituting this inverse covariance into Equation 112, we obtain

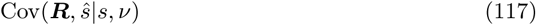

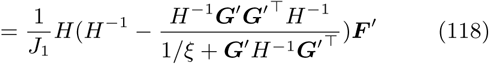

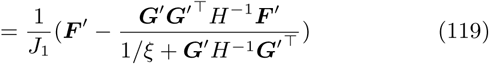

This last expression can be rewritten using elements of the Fisher information matrix, whose inverse bounds the covariance of any joint estimator of the signal and nuisance variables, 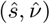:

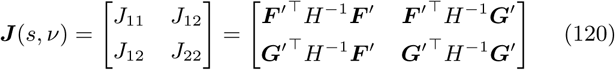

With these substitutions, we have

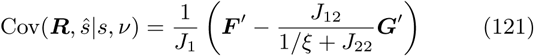

The denominator of Equation 111 involves the variance of the sufficient statistics,

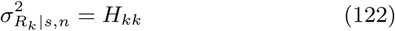

and the variance of the brain’s decoder,

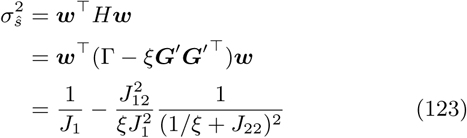

where we used the following results:

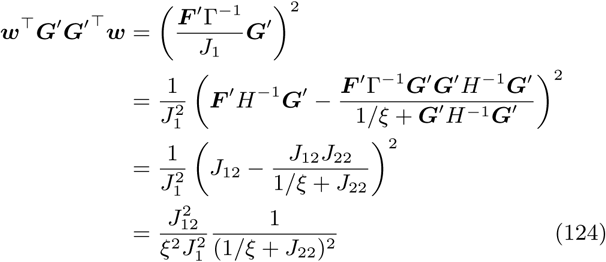

Combining the results from Equation 121, 123 and 122, we can compute Equation 111

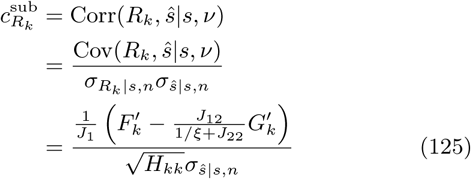

The optimal choice correlation when there is natural nuisance variation (Eq 7) is given by

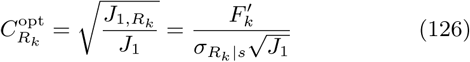

where 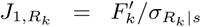 is the Fisher Information in *R*_*k*_ about *s* when there is natural nuisance variation, and 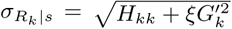 is the standard deviation of the statistic *R*_*k*_, again when there is natural nuisance variation.

The choice correlations for the same decoder differ under experimental conditions with and without nuisance variation: 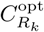 and 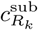. We find that the nuisance-conditioned choice correlations 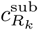 relate to the optimal nuisance-averaged choice correlations 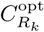 according to

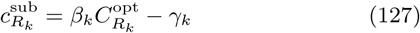

where we have defined the following constants:

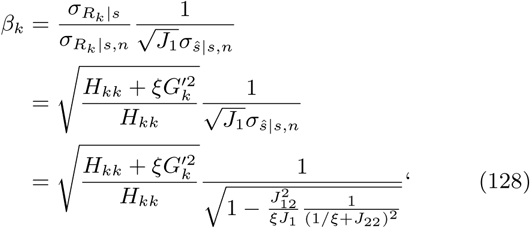

and

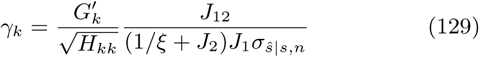

The slope *β*_*k*_ and offset *γ*_*k*_ of the relationship between these two types of choice correlations (Equation 127) depends on the amount of nuisance variation compared to internal noise and the suboptimality of the brain’s decoding strategy. When the signal and nuisance can be disentangled, that is, estimated nearly independently using the statistics ***R***(***r***), then *J*_12_ is small and the choice correlations driven purely by internal fluctuations closely match the optimal choice correlations in the presence of nuisance variation (Figure S2A). In contrast, when nuisance variations remain partialy confused with the signal, then *J*_12_ is large and the choice correlations for fixed nuisance variables may differ from the optimal pattern seen when allowing nuisance variables to change from trial to trial (Figure S2B).

For the simulations in Figure S2, we set the sufficient statistics to be linear ***R***(***r***) = ***r*** for simplicity. Neural responses were generated from a Gaussian distribution with a stimulus-dependent mean and identity covariance *H* = *I*: *p*(***r***|*s, ν*) = 𝒩(***F***′*s* + ***G***′*ν, I*). In Figure S2A, ***F***′ and ***G***′ are set to be orthogonal to ensure *J*_12_ = ***F***′^⊺^*H*^−1^***G***′ = 0. They are picked from the eigenvector of a symmetric matrix *A*^⊺^*A*, where *A* is a matrix whose elements are generated from uniform distribution bounded by 0 and 1. In Figure S2B, each element in ***F***′ and ***G***′ is drawn from a uniform distribution over the interval [0, 1]. We simulate 10000 responses of a population with *N* = 50 neurons. The stimulus is set to 0 and the nuisance is fixed to be 1. The brain’s decoder assumes a Gaussian prior over the nuisance variation with zero mean and variance *ξ* = 2. The decoding weights follow Equation 108, and the stimulus is estimated using Equation 109. Choice correlations in this fixed-nuisance experiment are computed by Equation 110 (vertical axis in Figure S2). The predicted optimal choice correlation is computed by Equation 126 (horizontal axis in Figure S2). In this setting, *β*_*k*_ ≈ 1 when *J*_12_ = 0.

**Figure S2:**
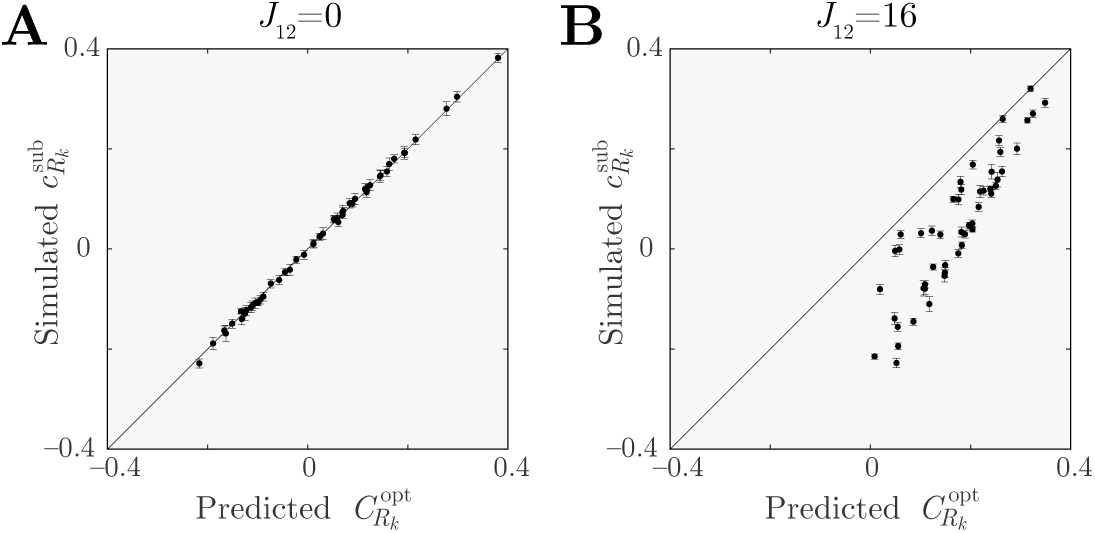
Comparing choice correlations caused by internal and external noise. (**A**) When estimates of nuisance variables are independent of estimates of task-relevant signals, the optimal choice correlations driven by internal noise, 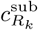, match the optimal pattern 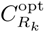 expected for optimal decoding under natural nuisance variation (Equation 7). (**B**) When the signal and nuisance variables remain confounded by an estimator and decoding is evaluated under different conditions than those for which it was optimized, then the choice correlations need not match this optimal prediction.

### S.5 Coarse discrimination and choice correlations

We now derive a relationship between nonlinear neural thresholds and nonlinear choice correlations for *coarse* binary discrimination tasks, choosing between stimulus *s*_+_ and *s*_−_. The main ideas are the same as for fine discrimination, but there are a few more subtleties involved when the statistical structure of the response depends on the stimulus.

We assume the brain decodes neural activity ***r*** as a linear weighted sum of nonlinear statistics ***R***(***r***), using weights given by linear regression as

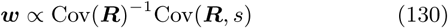

The latter factor reflects the signal strength,

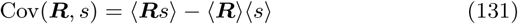

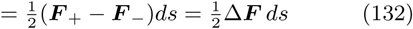

We assume that the two values *s*_±_ = *s*_0_ ±Δ*s* are equally probable, and notate the mean responses as ***F***_±_ = ***F*** (*s*_±_) = ⟨***R***|*s*_±_⟩. The factor Cov ***R*** includes covariability induced by both signal and noise. Using the law of total covariance, these contributions can be separated as

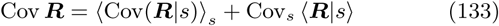

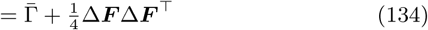

where the first term is the average noise covariance across the stimulus ensemble, 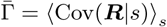, and the second term reflects variance along the signal direction. As for fine discrimination, noise variance along the signal direction has no influence on the optimal readout direction, since it cannot be removed. Using the Sherman-Morrison formula, we find that the decoder is

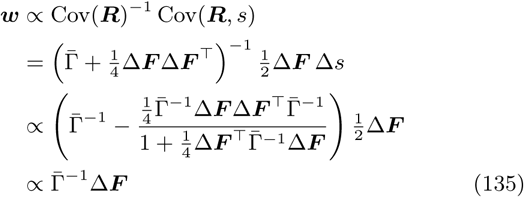

For unbiased decoding, the proportionality is given by 1*/*Δ***F***^⊺^Γ^−1^Δ***F***.

#### S.5.1 Average conditional choice correlations

The core desideratum for a measure of choice correlations is to isolate the non-stimulus fluctuations that correlate with choices. The typical way to ensure this is to measure correlations between neural responses and choices only when the stimulus is completely ambiguous, *i.e.* at the decision boundary. Other studies have sought to expand the range of stimuli that can be used for these correlations [38, 73]. Mathematically, we examine the statistical relationship between neural responses and choices that remains after *conditioning* on the stimulus, via *p*(***R***, *ŝ*|*s*). Here we quantify this relationship through a conditional covariance, Cov(***R***, *ŝ*|*s*). For coarse discrimination, the strength (and pattern) of this correlation may depend on the particular stimulus used. To account for this, we compute an average over possible stimuli, ⟨Cov(*R*_*k*_, *ŝ*|*s*)⟩_*s*_. If we normalize by root mean variances, we obtain

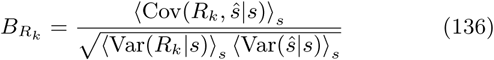

This nonlinear choice correlation can be rewritten as

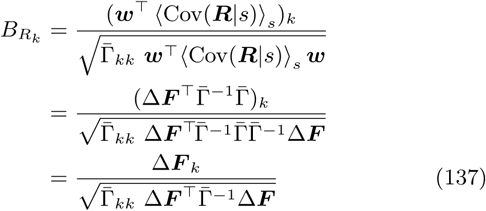

We recognize that this expression contains the ratio of sensitivities for the neural statistic *R*_*k*_ and the entire population ***r*** in coarse discrimination, 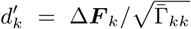 and 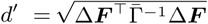. We therefore find the same result as for optimal fine discrimination (Eq 18):

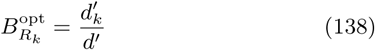

#### S.5.2 Signal estimation from total correlations

It is useful to express the discriminability through the total correlation between the responses and the stimulus,

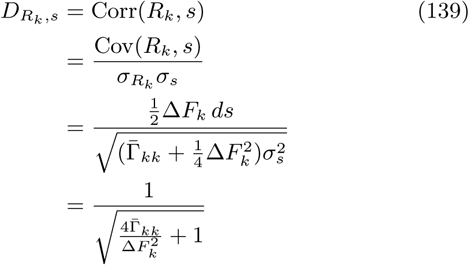

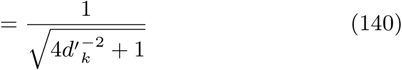

In these equations we used the fact that for binary discrimination, the standard deviation of the signal is related to the difference between the two possible signal values, 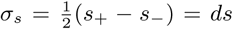. We can invert Eq 140 to find

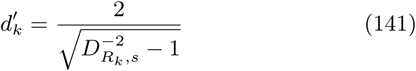

This dependence is plotted in Figure S3.

Similarly, we can express the behavioral discriminability *d*′ in terms of the correlation between the estimate and the stimulus, *D*_*ŝ,s*_ = Corr(*ŝ, s*):

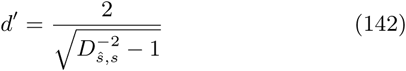

The relationship between discriminability *d*′ and total correlation *D* is linear when *D* is relatively small. Thus we can approximate the optimal nonlinear choice correlation as:

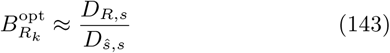

Here the total correlation is computed based on a continuous estimate *ŝ*. When the behavioral outcome is a binary choice, this relationship is more complicated. Section S.6.3 calculates the relationship between *D*_*ŝ*_ and 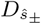 for one particular task.

### S.6 Orientation variance discrimination task

#### S.6.1 Coarse tasks: Continuous estimation versus binary discrimination

The experiment of Section 2.3 defines an orientation variance discrimination task in which the relevant statistics are quadratic functions of the orientation. The quadratic decoding model described in the main text could suffice for this problem. However, in our case the variances to be distinguished are quite different, such that the nuisance variation differs substantially between these two stimulus categories. As described in Methods 4.2.1, coarse tasks with stimulus-dependent variability generate a slightly different prediction compared to fine tasks (or coarse tasks with stimulus-independent variability).

**Figure S3:**
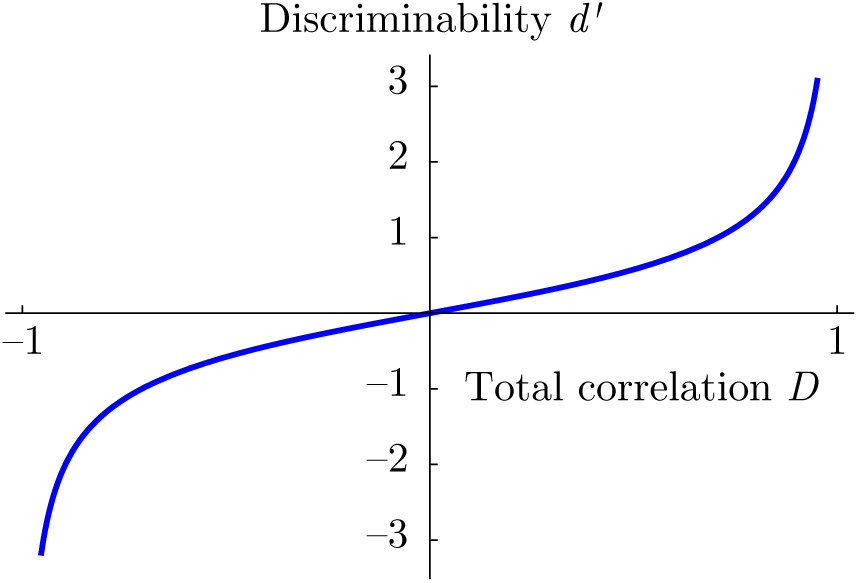
Stimulus discriminability *d*′ for a response variable *R*, versus total correlation between that variable and the stimulus, *D* = Corr (*R, s*), according to Eq 141.

Moreover, there are minor differences between the predictions for continuous estimation and binary discrimination, and these differences are more complicated for coarse tasks than fine ones. Here we describe in detail the somewhat lengthy computation of the ratio *ζ* between choice correlations for continuous quadratic estimation and binary quadratic decoding. For coarse discrimination, the ratio *ζ* will depend on the input statistics and threshold, but for fine discrimination *ζ* becomes a constant. Regardless, for our cases of interest these numbers are generally near 1.

We begin by assuming that the variance estimate is the square of the orientation estimate 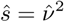, and a binary guess about the variance is given by 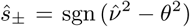 where *θ* is the animal’s orientation threshold. We assume that 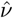 is an unbiased estimate of the orientation *ν*, so 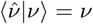. We denote one neuron’s mean response to the orientation by ⟨*r*|*ν*⟩ = *µ*(*ν*) which we approximate linearly as 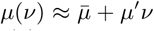 with 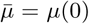. The mean behavioral choice is ⟨*ŝ*_±_|*s*⟩ = *m*_*s*_. Since the stimulus is binary, we will denote this mean with a subscript, ⟨*ŝ*_±_|*s*_+_⟩ = *m*_+_ or ⟨*ŝ*_±_|*s*_−_⟩ = *m*_−_.

The joint distribution 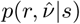 arises from both internal noise and nuisance variation, 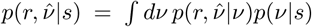. For a given orientation *ν*, the neural response *r* and orientation estimate 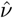 follow a bivariate normal distribution,

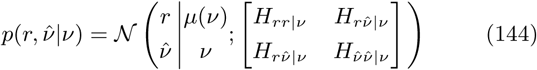

which summarizes all of the internal noise given the sensory input.

By design, the nuisance variable *ν* is normally distributed, *p*(*ν*|*s*) = 𝒩 (*ν*|0, *s*), so we can write the marginal distribution 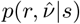 as

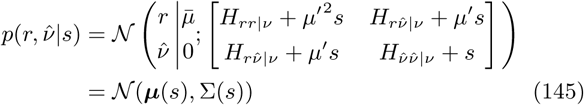

For now we suppress the explicit dependence on *s*.

The conditional covariance between the nonlinear statistic *R* and choice is

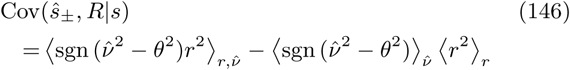

where *R* = *r*^2^ and we reiterate that we are suppressing the conditioning on *s*. The second moment is

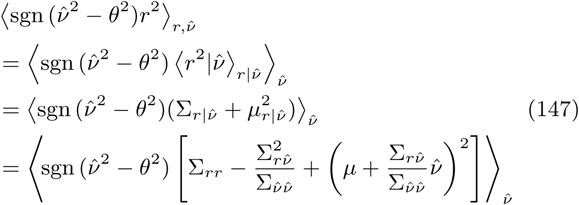

where we used the conditional distribution

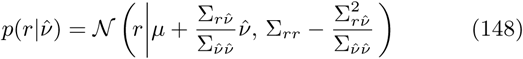

This can be written as

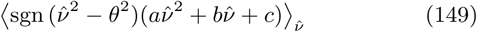

for coefficients

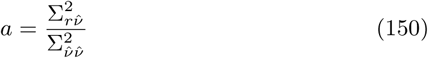

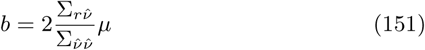

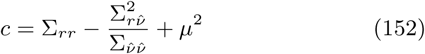

Note that this is an expectation over 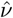 only. Such an expected value can be written as a sum of integrals:

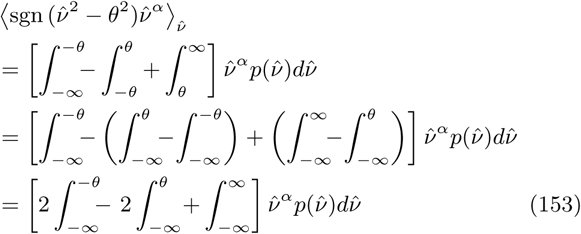

These integrals can be expressed in terms of error functions, where 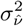 is the marginal variance for 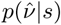:

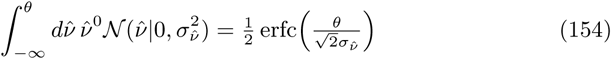

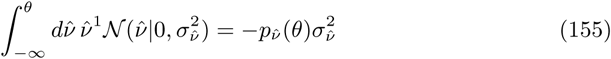

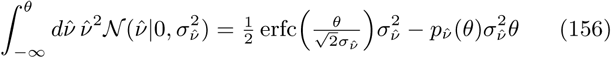

Note that 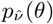 has units of [*ν*]^−1^, so units are consistent across these expressions.

Combining these with Eq 153 we obtain

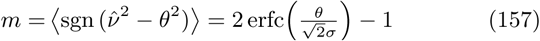

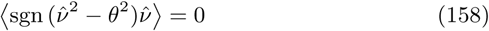

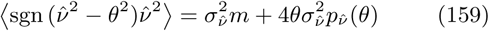

where we have used the identity erfc(−*x*) = 2 − erfc(*x*) and the symmetry 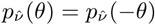. The first term, *m*, is the mean of *ŝ*_±_, and will appear several times in the equations below.

Returning to Eq 147, we have

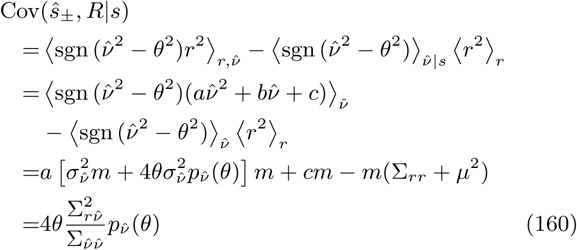

Note that all of the erfc terms have canceled.

Compare that to the corresponding covariance for continuous estimation,

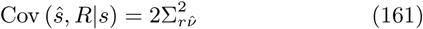

The conditional variance of a binary output *ŝ* = ±1 is simply

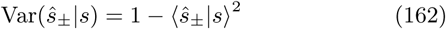

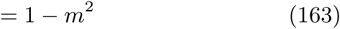

whereas, the variance for the continuous estimator is

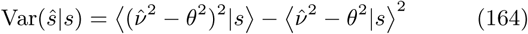

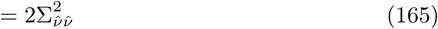

The variance of 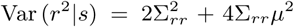, is the same whether the behavioral estimate is continuous or binary.

Our goal here is to compute the change in our measure of nonlinear choice correlation, namely,

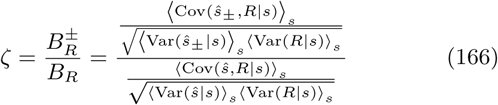

where the averages over *p*(*s*) = 1*/*2 include equal proportions of the binary stimuli *s*_+_ and *s*_−_. Substituting our calculations above, and reintroducing the dependencies on *s*, we find

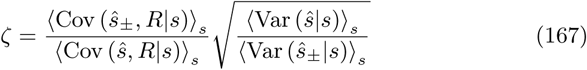

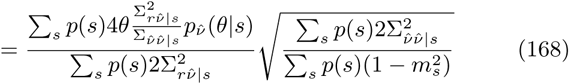

For tasks where the variability is dominated by external nuisance variables rather than by internal noise, i.e. *H* ≪ *s*(*µ*′, 1)(*µ*′, 1)^⊺^, we can approximate the covariances by 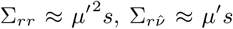, and 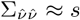. Substituting these approximations into the expression above, we obtain

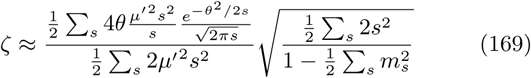

In our task conditions, 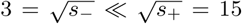, so some terms dominate in the sums. Moreover, we assume that the threshold *θ* lies far enough between 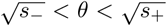 that 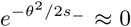 and 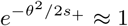. We then find

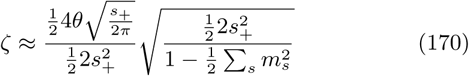

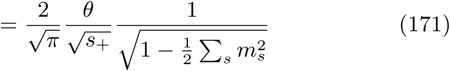

Empirically, we find that 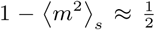 (Figure S4). In that case, we obtain

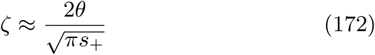

This expression is independent of the statistics of *r*. Therefore the same correction factor holds for cross-terms like *r*_*j*_*r*_*k*_, which can be expressed as linear combination of squares, 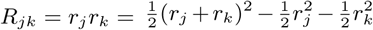. We use this correction factor *ζ* to adjust our predicted quadratic choice correlations in Figure 6.

To find the behavioral threshold *θ* for Eq 172, we used logistic regression of choice *ŝ*_±_ on the absolute value of the stimulus orientation, |*ν*|, and assign the threshold *θ* to be the orientation where the probability of both choices was equal.

**Figure S4:**
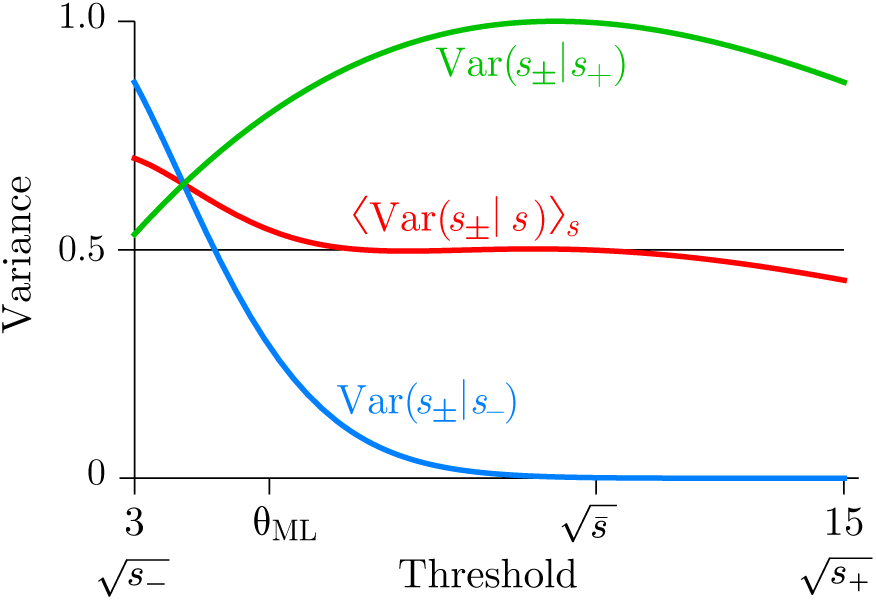
The average variance of *ŝ*_±_ conditioned on the stimulus *s* (red) is approximately 1/2 over a wide range of thresholds.

#### S.6.2 Fine tasks: Continuous estimation versus binary discrimination

For fine discrimination, the stimulus *s* is effectively constant, so we need not take averages.

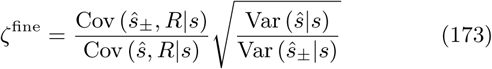

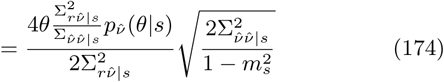

After several cancelations, and using the fact that for fine discrimination, *θ* = *s* ≈ *s*_+_ ≈ *s*_−_, we find

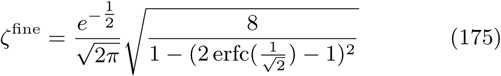

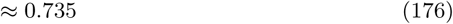

Observe that for fine discrimination, the ratio *ζ* is a constant, independent of the underlying statistics.

#### S.6.3 Total correlation for binary and continuous estimates

We showed in Methods 4.2.1 that the discriminability is related to the total correlation between signal and response. However, those relationships were based on continuous estimates of the binary stimulus. As above, when the behavioral choice is also binary, we can adjust the calculation slightly. Here we compare the total correlations for continuous and binary response, *D*_*ŝ,s*_ and 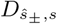.

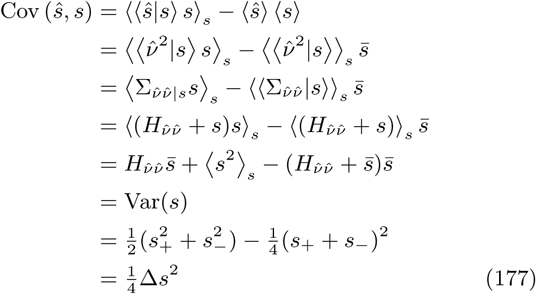

In contrast, the total covariance of *ŝ*_±_ is

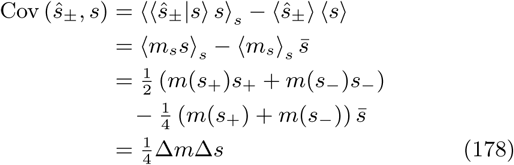

The total variance of *ŝ* is

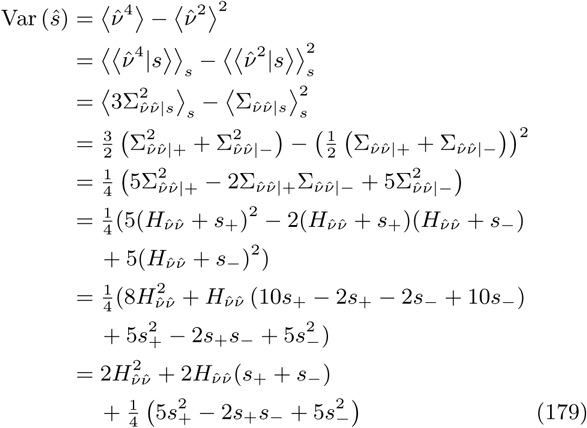

In the limit where the nuisance variability dominates the internal variability, and *s*_+_ ≫ *s*_−_, this simplifies to

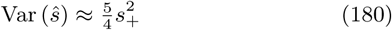

The total variance of *ŝ*_±_ is

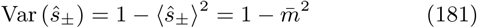

where 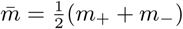.

Combining these computations, we see that ratio of total correlations for binary *ŝ*_±_ and continuous *ŝ* is

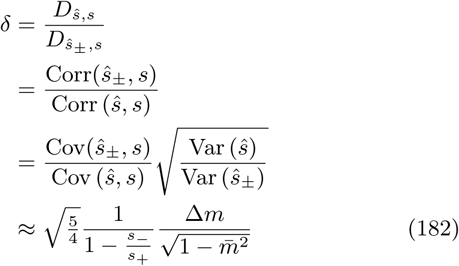

All of these quantities are measurable from data or are given by the task.

#### S.6.4 Optimal binary nonlinear coarse choice correlations

We can now combine our results above to create a prediction for optimal binary nonlinear coarse choice correlations. From Eq 143 and Eq 166, we have

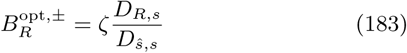

From Eq 182 we can adjust the

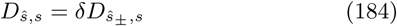

Combining these we have

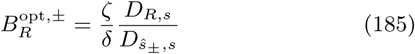

where *ζ* and *δ* are determined by experimentally measurable quantities. Their precise values depends on the monkey and the session, but the ratio is typically *ζ/δ* ≈ 0.62 ± 0.33. When plotting the data in Figure 6, we apply these corrections to each session before combining different sessions together.

